# Homologous mutations in β, embryonic, and perinatal muscle myosins have divergent effects on molecular power generation

**DOI:** 10.1101/2023.07.02.547385

**Authors:** Chao Liu, Anastasia Karabina, Artur Meller, Ayan Bhattacharjee, Colby J Agostino, Greg R Bowman, Kathleen M Ruppel, James A Spudich, Leslie A Leinwand

**Author notes:** Correspondence (L.A.L.).

## Abstract

Mutations at a highly conserved homologous residue in three closely related muscle myosins cause three distinct diseases involving muscle defects: R671C in β-cardiac myosin causes hypertrophic cardiomyopathy, R672C and R672H in embryonic skeletal myosin cause Freeman Sheldon syndrome, and R674Q in perinatal skeletal myosin causes trismus- pseudocamptodactyly syndrome. It is not known if their effects at the molecular level are similar to one another or correlate with disease phenotype and severity. To this end, we investigated the effects of the homologous mutations on key factors of molecular power production using recombinantly expressed human β, embryonic, and perinatal myosin subfragment-1. We found large effects in the developmental myosins, with the most dramatic in perinatal, but minimal effects in β myosin, and magnitude of changes correlated partially with clinical severity. The mutations in the developmental myosins dramatically decreased the step size and load-sensitive actin-detachment rate of single molecules measured by optical tweezers, in addition to decreasing ATPase cycle rate. In contrast, the only measured effect of R671C in β myosin was a larger step size. Our measurements of step size and bound times predicted velocities consistent with those measured in an in vitro motility assay. Finally, molecular dynamics simulations predicted that the arginine to cysteine mutation in embryonic, but not β, myosin may reduce pre-powerstroke lever arm priming and ADP pocket opening, providing a possible structural mechanism consistent with the experimental observations. This paper presents the first direct comparisons of homologous mutations in several different myosin isoforms, whose divergent functional effects are yet another testament to myosin’s highly allosteric nature.

## Introduction

Myosin myopathies refer to a group of skeletal and heart muscle diseases caused by mutations in sarcomeric myosin genes. There are ten such genes in mammals, and mutations in five of them have been linked to disease^1, 2^. These diseases are usually autosomal dominant and are largely the result of missense mutations. Some are relatively common, such as hypertrophic cardiomyopathy (HCM) that occurs at a frequency ∼1 in 500^3^ to 1 in 200 people^4^, while others are extremely rare such as Freeman Sheldon syndrome (FSS)^5^. They can be associated with a risk of sudden death as is the case with HCM, or not be associated with reduced lifespan but have profound effects on quality of life. A hallmark of the sarcomeric myosin gene family is the high sequence conservation among its members, which led to the early assumption that they were functionally redundant. Genetic inactivation of members of this gene family in mice has invalidated this assumption and demonstrated that the genes play distinct roles in heart and skeletal muscles^6–8^. Additionally, biochemical and biophysical analyses have described functional diversity among its members^9^.

In the current study, we evaluated the functional effects of mutations of the same highly conserved arginine residue in the motor domain of three different myosin genes that cause three distinct diseases (Figure 1). We hypothesized that all mutant motors would exhibit similarly altered biochemical and biophysical properties. However, we show here that each mutation results in distinct molecular phenotypes that partly correlate with clinical disease severity. The mutations are R671C in the MYH7 gene that causes HCM^10^, R672C and R672H in MYH3 that cause FSS^11^, and R674Q in MYH8 that causes trismus-pseudocamptodactyly syndrome (TPS)^12–14^.

**Figure 1.**
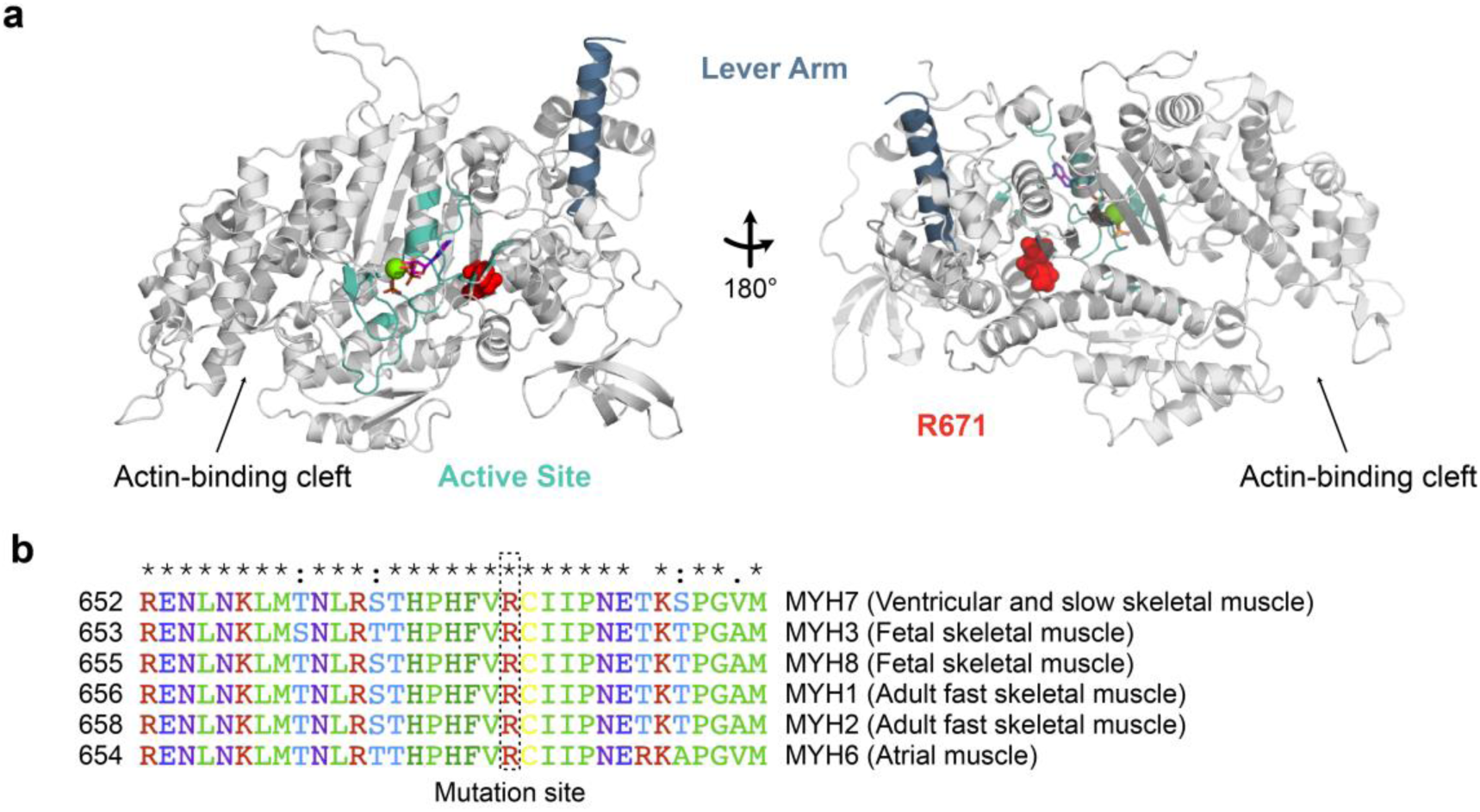
Mutations at a highly conserved location in the motor domain of different myosin isoforms cause distinct diseases. **a**. A homology model structure of β-cardiac myosin motor domain highlights that the arginine residue (R671) is located between the active site and lever arm. **b**. A multiple sequence alignment of different myosin-IIs expressed in various muscles shows that the mutation site is highly conserved. This paper studies homologous R671 mutations in β (MYH7), embryonic (MYH3), and perinatal (MYH8) myosins implicated in hypertrophic cardiomyopathy (HCM), Freeman-Sheldon syndrome (FSS), and trismus pseudocamptodactyly syndrome (TPS), respectively. Overall, sequence identity and similarity are respectively 79% and 89% between β and embryonic myosin, 81% and 90% between β and perinatal myosin, and 85% and 93% between embryonic and perinatal myosin.

The highly conserved R671 residue is located inside the myosin head between the active site and the lever arm (Figure 1a). It resides on the third β-strand and likely interacts with residues of the relay helix, SH1-SH2 domain, and adjacent β-strands^15^, regions that undergo major conformational changes during the myosin ATPase cycle^16^. Thus, mutations of this residue are expected to have consequential disruptions of myosin’s mechanoenzyme activity in all three isoforms.

The MYH7 gene is expressed in the heart and slow skeletal muscle in humans while the MYH3 and 8 genes, otherwise known as embryonic and perinatal, are expressed transiently in fetal developing skeletal muscle^17, 18^. The clinical features of HCM are a hypertrophied heart with hyperdynamic contractility and increased risks of atrial fibrillation, progressive heart failure, and sudden cardiac death^19^. Thousands of mutations across the genes encoding proteins of the cardiac sarcomere have been identified to cause HCM, with mutations in MYH7 being the second most common^19^. Recently, the FDA approved the use of a myosin inhibitor, Mavacamten, for symptomatic obstructive HCM^20^. FSS and TPS are congenital muscle contracture syndromes, specifically distal arthrogryposis types 2A and 7^21^. FSS is characterized by facial contractures (“whistling face”), strabismus, and contractures of the limbs, and is considered the most severe form of distal arthrogryposis^5, 22, 23^. Thus far, the only gene implicated in FSS is MYH3, which explains over 90% of cases^11^. Surgical interventions have been used to treat some of the clinical features of FSS, but no pharmacologic treatments exist. Trismus pseudocamptodactyly is caused by only one known mutation (R674Q) in MYH8 to date^14^. Patients have shortened muscles and tendons and cannot fully open their mouths. There is no treatment for this disease.

Until recently, studies to determine the functional effects of disease-causing mutations in striated muscle myosins relied on either patient biopsy samples or material from animal models. Analysis of patient samples is complicated by the presence of both wild type (WT) and mutant myosin of interest, as well as other myosin isoform(s). While no studies of cardiac biopsy material from patients with the MYH7 R671C mutation have been published, Racca et al. reported the effects of the MYH3 mutation R672C on muscle cells and myofibrils isolated from two patients with FSS^24^. They first confirmed that the embryonic heavy chain was expressed in adult skeletal muscle alongside the adult fast skeletal heavy chain encoded by MYH2, and that expression levels of the embryonic isoform were similar in cells from FSS patients and healthy controls. They then showed that the maximum specific force of the R672C-containing muscle cells was less than half of the controls, and that the mutant cells showed profoundly slower relaxation kinetics and incomplete relaxation. At the myofibril level, the initial slower phase of relaxation had a longer duration and slower rate, resulting in a greatly prolonged time to complete relaxation. Based on this and other data, they hypothesized that the R672C mutation may lead to very long-lived myosin cross-bridges with impaired detachment that result in the increased relaxation time and elevated passive tension seen at the myofibrillar and cellular levels. Another study of two other FSS mutations (Y583S and T178I) in *Drosophila* skinned muscle also proposed prolonged actin-myosin bound times as a key mechanism but, like Racca et al., lacked direct experimental evidence for this hypothesis^25^.

Analysis of the functional effects of disease-causing mutations using animal models is complicated by the fact that the mutation resides in a heterologous backbone with significant sequence differences from the human myosin motor in addition to the mutation of interest^26, 27^. Rao et al. created an animal model of FSS by introducing R672C to the *Drosophila* muscle myosin heavy chain gene and determining the effect on indirect flight muscle structure and function^28^. Flies heterozygous for the R672C mutation were unable to fly and exhibited severe myofibril disruption progressing to severe muscle degeneration. R672C mutant myosin isolated from the dissected indirect flight muscles of homozygous flies showed no change in the *k*_cat_ of the actin-activated ATPase but a 50% decrease in the gliding velocity of actin in an in vitro motility assay. Homology modeling suggested that the mutation disrupts communication between the nucleotide-binding site and the lever arm.

Finally, using a recombinant mammalian expression system for producing mutant and wild type human myosins^9, 29, 30^ that overcomes the limitations of the systems described above, Walklate et al. studied the effects of the R672C and R672H mutations on homogeneously purified populations of the human embryonic myosin motor domain (subfragment 1, S1)^15^. They found that each mutation resulted in a decrease in the *k*_cat_ of the actin-activated ATPase and an increase in the apparent affinity for actin (decreased *K*_m_), with the histidine substitution having a greater effect. In transient kinetic studies, the mutant proteins showed significantly decreased ATP hydrolysis rates, increased affinity for actin in the presence of ADP, and significantly decreased ATP-induced detachment rates from actin. R672H also showed enhanced affinity for ADP. Based on a loss of amplitude of the intrinsic tryptophan fluorescence signal in the mutant proteins, the authors hypothesized that the mutations disrupted a key interaction between R672 and F490 on the relay helix to result in the relay helix not moving properly during the recovery stroke.

In this report, we extend the analysis of the MYH3 R672C and R672H mutants to include characterization of their biophysical properties at the single molecule and ensemble levels. We then compare these mutant proteins to β and perinatal myosins harboring mutations at the homologous arginine residue. Surprisingly, despite the high sequence conservation amongst the different myosin isoforms in the region of the arginine residue, each mutant myosin isoform displayed a distinct biochemical and biophysical phenotype, albeit similarities existed between the developmental mutants. To our knowledge, this is the first report of the biochemical and biophysical effects of homologous mutations in several different myosin isoforms.

## Results

In order to understand how mutations in the same residue of three closely related MYH genes (Figure 1) cause three diseases with very different phenotypes, we measured the biochemical and biophysical properties of recombinant human myosin motor domains (S1 including both the essential and regulatory myosin light chain binding domains) of MYH7 (β, “βS1”), MYH3 (embryonic, “EmbS1”) and MYH8 (perinatal, “PeriS1”). Constructs consisted of S1 recombinantly expressed in mouse myoblast C2C12 cells^9^, bound by coexpressed human essential light chain^31^ and endogenous mouse regulatory light chain, and with a C-terminal 8-amino acid tag for purification and surface attachment purposes (Supplementary Figure S1). Each wild type (WT) myosin was compared to myosins bearing the following mutations: βS1-R671C, EmbS1-R672C, EmbS1-R672H, and PeriS1-R674Q. Given the sequence conservation of this region of the molecule (FIgure 1), one may hypothesize that the properties of the mutant motors would be similar to each other and that the diverse disease phenotypes would be the result of time and tissue distribution differences in the expression of these three myosins. However, as the myosin molecule is highly allosteric, it is not entirely unexpected that our results showed divergent effects of the mutations in the different isoforms. Further investigation of this allosteric mechanism using molecular dynamics simulations to compare the same mutation in βS1 (R671C) and EmbS1 (R672C) suggests that differences in pre-powerstroke conformations may account for the experimentally observed differences in motor function.

### Homologous mutations had differential effects on developmental and β myosin ATPase activity

Actin-activated ATP hydrolysis rates were measured in an NADH-coupled solution based assay, and the data were fit to the Michaelis-Menten model for enzyme kinetics to determine *k*_cat_ and *K*_m_ of the S1 motors. The WT developmental skeletal motors demonstrated a higher ATPase turnover than WT β myosin (Figure 2a-e, Table 1) consistent with previous comparisons of human myosin ATPase rates^9, 15, 32^. The EmbS1 and PeriS1 *k*_cat_ = 8.8 ± 0.6 s^-1^ and 11.0 ± 4.0 s^-1^, respectively, versus *k*_cat_ = 4.2 ± 0.8 s^-1^ for βS1 (mean ± s.e.m. are reported for all measured values throughout this paper. *p*-values of pairwise comparisons are given in Supplemental Table 1). The mutations considerably slowed embryonic and perinatal ATPase rates (*k*_cat_ = 6.5 ± 0.4 s^-1^ for R672C EmbS1, 4.3 ± 0.3 s^-1^ for R672H EmbS1, 2.4 ± 0.5 s^-1^ for R674Q PeriS1). However, despite a very large difference in mean values, the difference between *k*_cat_ of WT and R674Q PeriS1 was not significant due to a wide spread of the WT PeriS1 data. This was not surprising because the high *K*_m_ values of WT PeriS1 prevented accurate determination of *k*_cat_ as the plateau phase was not well defined in this case. The R671C mutation did not change the ATPase turnover of the β motor (*k*_cat_ = 4.9 ± 1.0 s^-1^). This lack of a major effect on ATPase rate is a common trend seen with many HCM-causing mutations in β-cardiac myosin^33^.

**Figure 2.**
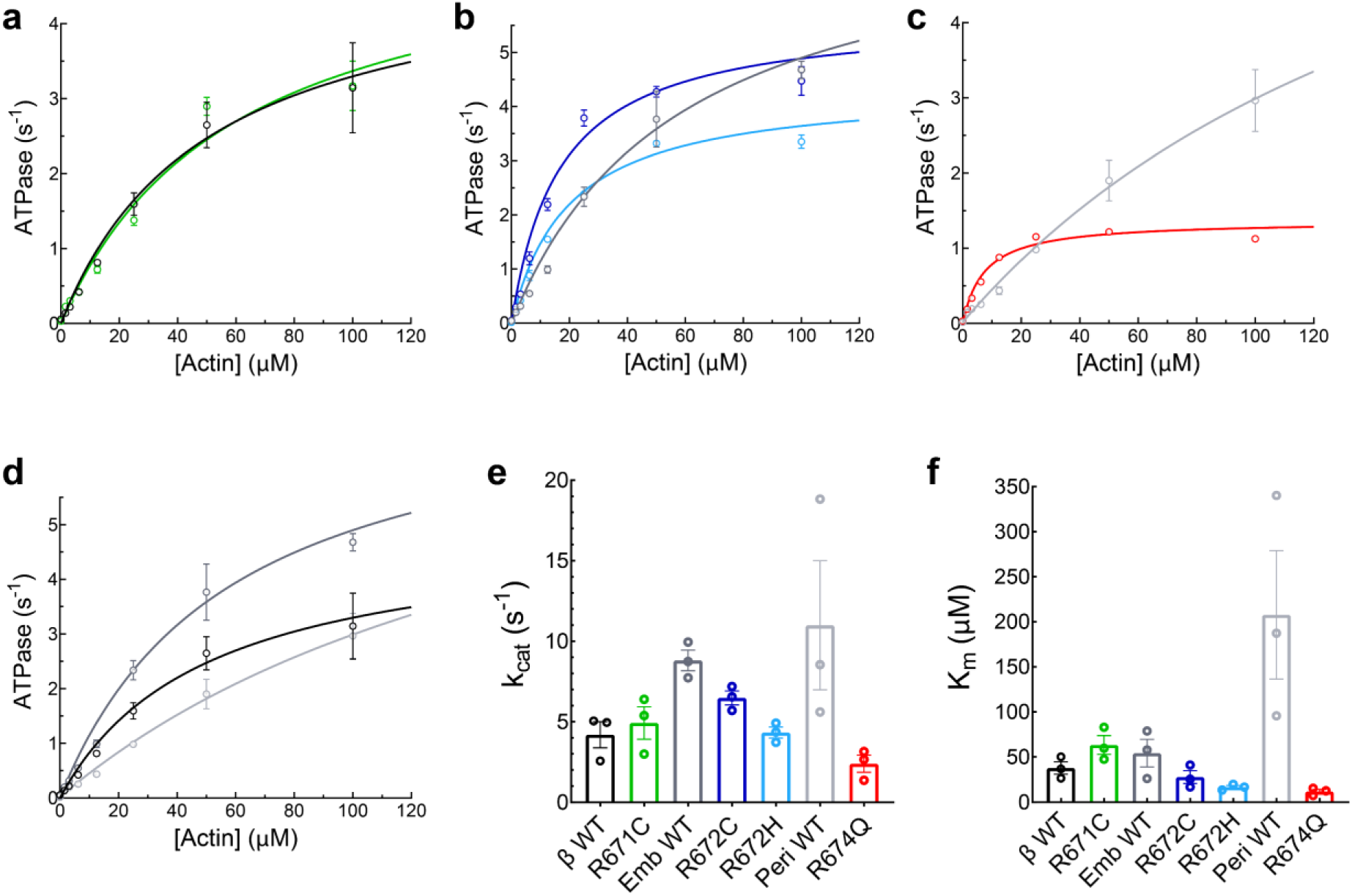
Steady-state actin-activated ATPase rates of myosins measured by solution absorbance. A representative dataset is shown for a single biological replicate of β-WT and R671C (**a**), EmbWT, R672C, and R672H (**b**), PeriWT and R674Q (**c**), with all three WT isoforms shown in (**d**) where each data point represents the average of three technical replicates at each actin concentration, and error bars represent standard deviations. The data is fit to the Michaelis- Menten equation (solid line) to determine *k*_cat_ and *K*_m_ for each myosin. Three independent experiments, each with their own curve fit, were performed at 30°C for each protein. Average values of *k*_cat_ (**e**) and *K*_m_ (**f**) are plotted as a bar graph with s.e.m., where data points represent the individual measurements of each independent experiment.

**Table 1.** Biochemical and biophysical parameters of myosins. ATPase (Michaelis-Menten rate *k*_cat_ and constant *K*_m_, measured at 30 °C), in vitro motility (velocities *V*_23°C_ and *V*_30°C_ and temperature coefficient *Q*_10_), and single molecule optical trapping (actin detachment rate at zero force *k*_0_, load sensitivity parameter δ, and step size *d*, measured at 23 °C). Duty ratio at zero force (*dr*_0_) and 30 °C is calculated by *k*_cat_/*k*_0_, where *k*_0_ at 30 °C is obtained by correcting the measured 23 °C *k*_0_ with *Q*_10_. Values of measured parameters are mean ± s.e.m. Uncertainties on calculated parameters (*Q*_10_ and *dr*_0_) are propagated errors. Statistical significance of relevant pairwise comparisons are given in Supplementary Table 1.

| | | $k_{\text{cat}}$ (s<sup>-1</sup>) | $K_m$ (μM) | $V_{23^\circ\text{C}}$ (nm/s) | $V_{30^\circ\text{C}}$ (nm/s) | $Q_{10}$ | $k_0$ (s<sup>-1</sup>) | $\delta$ (nm) | $d$ (nm) | $dr_0$ |
| --- | --- | --- | --- | --- | --- | --- | --- | --- | --- | --- |
| $\beta$ | WT | 4.2 ± 0.8 | 38 ± 7 | 837 ± 5 | 1727 ± 15 | 2.81 ± 0.04 | 174 ± 6 | 0.72 ± 0.05 | 4.7 ± 0.4 | 0.012 ± 0.002 |
| | R671C | 4.9 ± 1.0 | 63 ± 10 | 988 ± 15 | 2080 ± 39 | 2.90 ± 0.10 | 169 ± 7 | 0.73 ± 0.03 | 7.0 ± 0.5 | 0.014 ± 0.003 |
| Embryonic | WT | 8.8 ± 0.6 | 54 ± 15 | 1473 ± 28 | 3103 ± 40 | 2.90 ± 0.10 | 150 ± 14 | 0.95 ± 0.06 | 5.5 ± 0.3 | 0.028 ± 0.003 |
| | R672C | 6.5 ± 0.4 | 28 ± 7 | 337 ± 5 | 759 ± 3 | 3.19 ± 0.07 | 74 ± 5 | 0.37 ± 0.06 | 2.1 ± 0.3 | 0.039 ± 0.004 |
| | R672H | 4.3 ± 0.3 | 17 ± 2 | 170 ± 2 | 370 ± 1.3 | 3.04 ± 0.05 | 19 ± 2 | 0.72 ± 0.05 | 0.7 ± 0.3 | 0.102 ± 0.012 |
| Perinatal | WT | 11.0 ± 4.0 | 208 ± 71 | 1920 ± 35 | 4065 ± 20 | 2.92 ± 0.08 | 270 ± 15 | 0.76 ± 0.05 | 7.7 ± 0.5 | 0.019 ± 0.007 |
| | R674Q | 2.4 ± 0.5 | 12 ± 2 | 65 ± 3 | 130 ± 2 | 2.69 ± 0.19 | 6.2 ± 0.4 | 1.24 ± 0.09 | 0.7 ± 0.2 | 0.195 ± 0.043 |

The apparent affinity of myosin for actin, described by *K*_m_, was not significantly changed across wild type myosins. However, the *K*_m_ was differentially affected by the homologous mutations in the developmental versus β isoforms (Figure 2f, Table 1). The homologous mutations increased the actin affinity of the developmental myosins but slightly decreased the actin affinity for β myosin. βS1-R671C modestly increased the *K*_m_ of the motor by 66%, from 38 ± 7 uM (β-WT) to 63 ± 10 uM. R674Q-PeriS1 showed the most dramatic effect on *K*_m_ of all four homologous mutations with a 17-fold increase in actin affinity (PeriS1 *K*_m_ = 208 ± 71 uM, R674Q *K*_m_ = 12 ± 2 uM), whereas the EmbS1 mutations resulted in a ∼2 - 3-fold increase in actin affinity (WT *K*_m_ = 54 ± 15 uM, R672C *K*_m_ = 28 ± 7 uM, R672H *K*_m_ = 17 ± 2 uM).

Two previous studies, one using *Drosophila* myosin^34^ and the other using the same construct as our present paper^15^, reported the same trends in steady-state ATPase between EmbS1 and the R672C/H mutations, albeit the latter observed much more pronounced effects on *k*_cat_ and *K*_m_ than seen here. However, that study was performed in 0 mM KCl while the assays performed here contained 50 mM KCl to more closely mimic physiological conditions. It is known that the actin-myosin interaction is weakened with increasing salt concentrations^35^, which may explain the difference in the magnitude of the effects observed. Overall, the homologous mutations show a much greater effect on the enzymatic ATPase properties of the developmental skeletal myosin isoforms than on β myosin.

### Homologous mutations had opposite effects on motility velocity

The WT developmental myosins had significantly higher actin-sliding velocity in an unloaded motility assay than β myosin, as expected of the skeletal vs. cardiac isoforms (Figure 3, Table 1, Supplementary Movies 1,3,6). EmbS1 had velocity *V* = 1473 ± 28 nm/s at 23 °C, and PeriS1 had velocity *V* = 1920 ± 35 nm/s. All comparisons of measured parameters hereon are significant unless stated otherwise (e.g. “similar”, “not changed”), and *p*-values are given in Supplementary Table 1. Previously published velocities of the developmental myosins purified from human fetal tissues^36^ or from recombinant expression in C2C12 cells^9^ are somewhat lower than the values above, likely due in part to their failure to copurify the light chains^9^. βS1 had velocity *V* = 837 ± 5 nm/s, consistent with values previously published for human β-cardiac myosin^37–44^.

**Figure 3.**
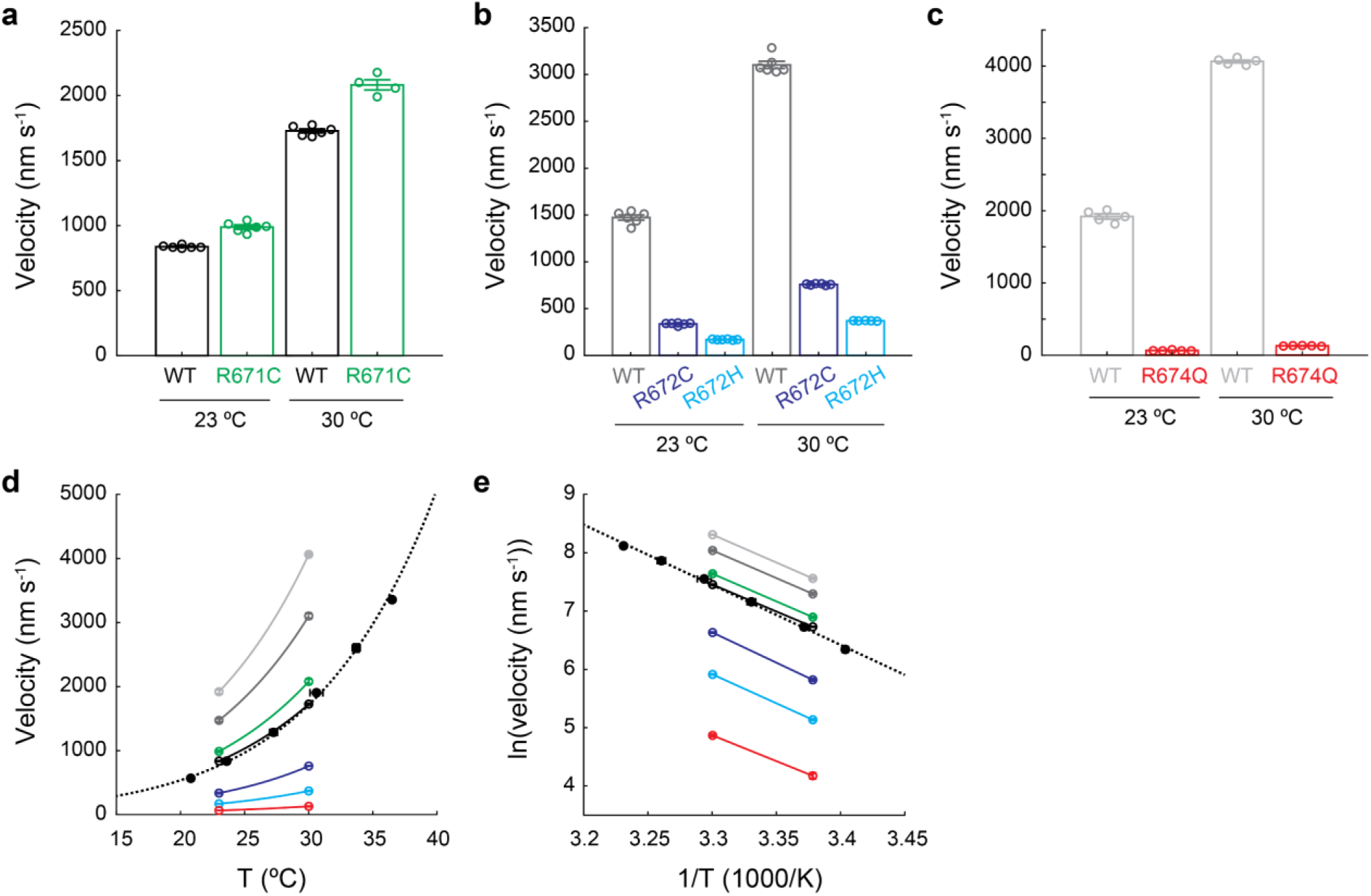
Actin sliding velocities of myosins measured by the in vitro motility assay. **a-c.** Velocities of β S1 WT and R671C (**a**), embryonic S1 WT, R672C, and R672H (**b**), and perinatal S1 WT and R674Q (**c**), each at 23 °C and 30 °C. Reported velocities are “Mean velocities filtered” (see *Methods* and Supplementary Figures 2-4 for detailed analysis and additional parameters obtained from motility data). Each data point represents one independent experiment. Error bars (most of which are smaller than the lines’ thickness) over the bar graph represent mean ± s.e.m. All relevant pairwise comparisons are statistically significant (Supplementary Table 1). **d**. Velocities of each myosin are plotted as a function of temperature. The pairs of data points at 23°C and 30 °C (open circles) are taken from **a-c**. The data points spanning ∼21 °C to 37 °C (black solid circles) are of wild type human β-cardiac short-S1 (residues 1-808) as reported in (Liu C, Ruppel KM, Spudich JA. (2023). Motility assay to probe the calcium sensitivity of myosin and regulated thin filaments. In M Regnier and M Childers (Eds.) *Methods in Molecular Biology: Familial Cardiomyopathies*. New York, NY: Springer Nature). Error bars represent mean ± s.e.m. Solid and dashed lines represent fits to the Arrhenius equation. **e.** Data in **d** displayed as the logarithm of velocity plotted against reciprocal of temperature. All values of velocities and temperature coefficients *Q*_10_’s (from Arrhenius equation fit) are given in Table 1. The color scheme introduced here is maintained throughout the paper: black, dark gray, and light gray represent the wild types of β, embryonic, and perinatal S1, respectively; green, blue, cyan, and red represent the various mutants as shown. Motility experiments were done using 2 mM ATP.

The homologous mutations dramatically decreased the actin-sliding velocity of the developmental myosins but slightly increased the velocity of β myosin. R671C increased the velocity of βS1 by 18% (*V* = 988 ± 15 nm/s at 23 °C) (Figure 3a, Table 1, Supplementary Movie 2). In contrast, R672C and R672H decreased the velocity of EmbS1 by 77% and 88%, respectively (R672C *V* = 337 ± 5 nm/s, R672H *V* = 170 ± 2 nm/s at 23 °C) (Figure 3b, Table 1, Supplementary Movies 4,5). A previous paper had reported a 50% reduction in motility velocity by R672C in myosin purified from *Drosophila*^28^. R674Q decreased the velocity of PeriS1 by 97% (*V* = 65 ± 3 nm/s at 23 °C) (Figure 3c, Table 1, Supplementary Movie 7).

Several mutations in β-cardiac myosin have been previously reported to cause poor protein expression and possibly improperly folded heads, or “deadheads”, that contribute in part to reduced motility with higher percentages of stuck actin filaments (eg. the dilated cardiomyopathy mutation S532P^38, 42^ and the hypertrophic cardiomyopathy mutation P710R^44^) (see also supplementary discussion in Vander Roest *PNAS* 2021). In the present work, however, protein expression and the presence of deadheads cannot explain the observed effects on motility velocities for any of the mutations. R671C βS1 had higher velocity than WT despite having lower protein expression (see note in Methods) and higher percentage of stuck actin filaments (Supplementary Figure S4c). R672C, R672H, and R674Q all had very low stuck percentages (∼5%) equal or lower than their respective WT proteins (Supplementary Figure S4c). The smoothness and quality of actin sliding are also evident in the motility movies (Supplementary Movies 1-14). Thus, the observed significant changes in velocities are direct effects of the mutations on motor function rather than due to altered expression or number of deadheads.

To determine whether the mutations alter the temperature sensitivity of myosins and to more closely mimic the physiological temperature, we measured motility velocity at 30 °C in addition to 23 °C. WT βS1 had temperature coefficient *Q*_10_ = 2.81 ± 0.04 (error is propagated from calculation of *Q*_10_) (Figure 3de, Table 1), consistent with the value measured for human β-cardiac sS1-AC, a construct consisting of the first 808 residues of the myosin heavy chain along with the essential light chain (*Q*_10_ = 3.11 ± 0.09) (Liu C, Ruppel KM, Spudich JA. (2023). Motility assay to probe the calcium sensitivity of myosin and regulated thin filaments. In M Regnier and M Childers (Eds.) *Methods in Molecular Biology: Familial Cardiomyopathies*. New York, NY: Springer Nature). WT EmbS1 and WT PeriS1 had similar temperature sensitivities to βS1 (EmbS1 *Q*_10_ = 2.90 ± 0.10, PeriS1 *Q*_10_ = 2.92 ± 0.08) (Figure 3de, Table 1). Each homologous mutation had the same relative effect on their respective WT myosin at both 23 °C and 30 °C (Figure 3, Table 1). While R672C had a small increase (*Q*_10_ = 3.19 ± 0.07), the temperature sensitivity was largely invariant in the myosins we studied, independent of isoforms or mutations.

### Homologous mutations greatly decreased the load-dependent actin detachment rate of single molecules of embryonic and perinatal myosin

Motor function of single myosin molecules was assessed by optical trapping using harmonic force spectroscopy (HFS). In this technique, binding events between a single myosin molecule and an actin filament are measured under different load forces. The sample stage oscillates harmonically so that by the randomness of where myosin initially binds actin, a range of mean forces are automatically applied over the course of many binding events^45^ (Figure 4a). This oscillation at high frequency (200 Hz) also enables the detection of cardiac and skeletal myosins’ short millisecond-length binding events under physiological (2 mM) ATP conditions. HFS has been used to measure the load-dependent kinetics and step size of β-cardiac myosin and the effects of numerous cardiomyopathy-causing mutations and myosin activators and inhibitors^42, 44, 46^.

**Figure 4.**
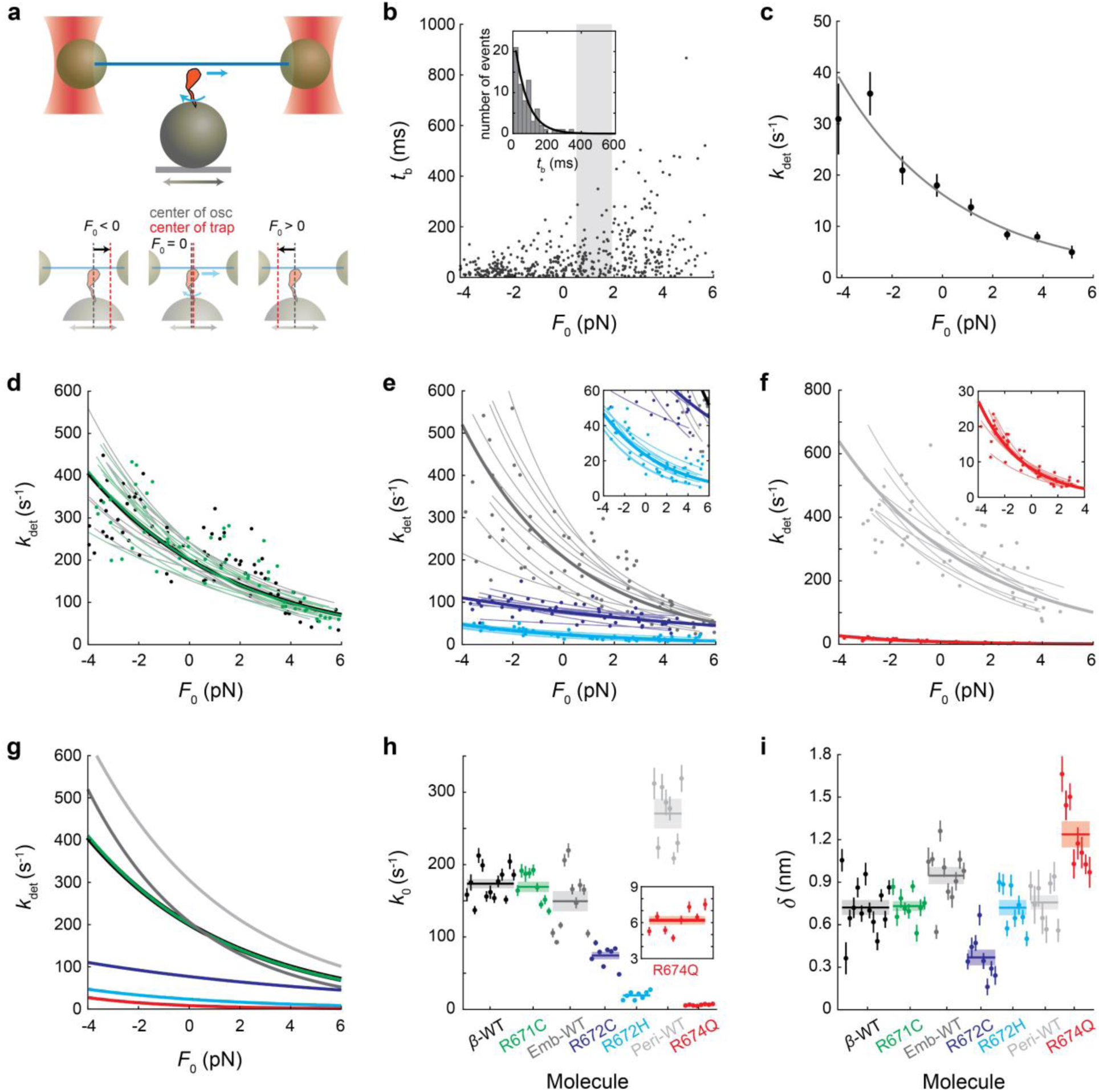
Single-molecule load-dependent actin-detachment kinetics of myosins measured by optical trapping. **a.** Setup of the three-bead system in the optical trap (top). In harmonic force spectroscopy (HFS), the stage upon which myosin sits oscillates so that a sinusoidal external force with mean *F*_0_ is applied to myosin upon actin binding. Depending on the randomness of when attachment occurs, binding events have different *F*_0_’s (bottom). *F*_0_ > 0 represents resistive force, opposite of the direction of stroke, and *F*_0_ < 0 represents assistive force, in the same direction of the stroke. Here, the direction of the lever arm swing and the resulting translation of the actin filament are indicated by the blue curved and straight arrows, respectively. See also Supplementary Figure 5 for illustration of the method of HFS to detect a binding event. **b.** All events for one example molecule. Time bound *t*_b_ of each event is plotted against *F*_0_. Events binned by force (example bin shown as shaded rectangle) have exponentially distributed bound times, from which the detachment rate at that force is determined by maximum likelihood estimation (MLE) (inset). **c.** Detachment rates *k*_det_ determined across all force bins for the example molecule from **b**. Error bars represent fitting errors from MLE. The force dependent Arrhenius equation with harmonic force correction (Equation 1) is fitted to the data and yields rate at zero load *k*_0_ = 12.5 ± 0.8 s^-1^ and force sensitivity δ = 0.88 ± 0.08 nm for this example R672H molecule. Errors on *k*_0_ and δ of one molecule represent fitting errors. See Supplementary Figure 6 for an example molecule of each protein. **d-f.** Load dependent detachment curves of all molecules of β S1 WT and R671C (**d**), embryonic S1 WT, R672C, and R672H (**e**), and perinatal S1 WT and R674Q (**f**). Each thin line represents one molecule. Data points are as described in **b** and **c** but displayed without error bars for clarity. Thick curves represent the mean across molecules, also plotted in **g** for comparison. **h-i.** *k*_0_ (**h**) and δ (**i**) for each molecule, each pair of parameters corresponding to one thin line in **d-f**. Error bars represent fitting errors. Horizontal thick lines represent means of *k*_0_ and δ across molecules, which define the thick curves in **d-g**. Shaded rectangles represent s.e.m. All values are given in Table 1. All relevant pairwise comparisons are statistically significant except for the following: βS1 WT vs R671C *k*_0_ and δ, βS1 WT vs EmbS1 WT *k*_0_, and βS1 WT and PeriS1 WT δ (Supplementary Table 1). Optical trap experiments were done at 23 °C using 2 mM ATP.

Analysis of HFS data is briefly outlined as follows (see Methods, Sung *Nat. Commun.* 2015^45^, and Liu *NSMB* 2018^42^ for additional details). Binding events are automatically detected from the recorded time series of the trapped beads’ positions based on a simultaneous increase in oscillation amplitude and decrease in phase relative to the stage position (Supplementary Figure S5). Both criteria occur due to strong coupling between the stage and the trapped beads when myosin binds the actin filament. The duration and mean load force are determined from each event (Figure 4b). Durations of events within a force bin are exponentially distributed with a characteristic rate (Figure 4b inset), the rate of detachment from actin at that load force.

Myosin’s rate of detachment from actin (*k*_det_) is an exponential function of the mean load force (*F*_0_) (Figure 4c):

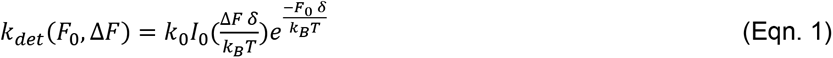

where *k*_0_ is the rate at zero load, δ is the distance to the transition state of the rate-limiting step in the bound state (a measure of force sensitivity), *k*_B_ is the Boltzmann constant, *T* is temperature, and *I*_0_ is the zeroth-order modified Bessel function of the first kind (to correct for the harmonic force with amplitude Δ*F*)^45^. In cardiac and skeletal myosins, ADP release is the rate-limiting step for detachment from actin at saturating (2 mM) ATP concentrations^47, 48^, and this step is sensitive to load force^49, 50^. Thus, the detachment rate *k*_0_ and its load sensitivity δ determined by HFS correspond to the rate of ADP release and its load sensitivity, respectively^50^.

WT β and embryonic myosins had similar actin detachment rates (βS1 *k*_0_ = 174 ± 6 s^-1^, EmbS1 *k*_0_ = 150 ± 14 s^-1^; *p* = 0.14) (Figure 4degh, Supplementary Figure S6ac, Table 1), comparable to values previously reported for β-cardiac myosin^45 42, 44, 51, 52^. WT perinatal myosin had significantly faster kinetics (*k*_0_ = 270 ± 15 s^-1^) (Figure 4f-h, Supplementary Figure S6f, Table 1). The force sensitivity of the actin detachment rate was similar in WT βS1 (δ = 0.72 ± 0.05 nm) and PeriS1 (δ = 0.76 ± 0.05 nm) (*p* = 0.63), while that of EmbS1 was slightly higher (δ = 0.95 ± 0.06 nm) (Figure 4i, Table 1). Measurement of PeriS1’s very short binding events (∼5-30 ms) at saturating ATP concentration under load was enabled by harmonic force spectroscopy’s use of stage oscillations; methods that rely on reduction in Brownian motion upon strong myosin-actin interactions can only accurately detect events of longer durations^51, 53–55^ and thus would likely fail to measure the detachment kinetics of faster myosins like PeriS1. While the 200 Hz stage oscillations used in our experiments set the shortest detectable events at 5 ms or one period of oscillation, increasing the oscillation frequency, within practical constraints like stage capabilities, would allow detection of even shorter binding events in other myosin isoforms that have even faster kinetics.

R671C did not change the actin detachment rate of βS1 or its force sensitivity (*k*_0_ = 169 ± 7 s^-1^, *p* = 0.65; δ = 0.73 ± 0.03 nm, *p* = 0.87) (Figure 4d, g-i, Table 1). WT and R671C βS1 molecules bound actin for ∼6 ms on average under zero load in the exponential distribution of bound times; assistive and resistive load forces up to 4 pN altered the average bound time to ∼3 ms and ∼15 ms, respectively (Supplementary Figure S6ab).

Both R672C and R672H significantly decreased the detachment rate of EmbS1 by 51% and 87% and its force sensitivity by 61% and 24%, respectively (R672C: *k*_0_ = 74 ± 5 s^-1^, δ = 0.37 ± 0.06 nm; R672H: *k*_0_ = 19 ± 2 s^-1^, δ = 0.72 ± 0.05 nm) (Figure 4e, g-i, Table 1). To put in perspective these detachment rates, WT EmbS1 molecules bound actin for ∼7 ms on average under zero load; assistive and resistive forces up to 4 pN altered the average bound time to ∼2 ms and ∼10 ms, respectively (Supplementary Figure S6c). R672C molecules bound actin longer than WT, for ∼10 ms, ∼14 ms, and ∼20 ms on average under -4 pN (assistive), 0 pN, and +4 pN (resistive) loads, respectively. Its much reduced force sensitivity is apparent through both the more symmetrical shape of the plot of bound times vs. load force and the shallower slope of the *k*_det_ vs load force relation (Supplementary Figure S6d). R672H molecules bound actin much longer than both WT and R672C for ∼30 ms, ∼50 ms, and ∼140 ms on average under -4 pN, 0 pN, and +4 pN loads, respectively. Its greater force sensitivity when compared with R672C is apparent through the more asymmetrical shape in both aforementioned plots (Supplementary Figure S6e). These single molecule results of slower detachment rates agree in trend with previously reported stopped-flow solution measurements of the ADP release and ATP binding rates: both R672C and R672H had slower rates than WT EmbS1, with the latter having a larger effect^15^.

R674Q dramatically decreased the detachment rate of PeriS1 by 98% while increasing its force sensitivity by 63% (*k*_0_ = 6.2 ± 0.4 s^-1^; δ = 1.24 ± 0.09 nm) (Figure 4f-i, Table 1). While WT PeriS1 molecules bound actin for ∼2-10 ms on average under different load forces, R674Q molecules stayed bound for ∼50 to hundreds of milliseconds (Supplementary Figure S6fg).

In summary, we measured the actin detachment kinetics of single myosin molecules under load using HFS optical trapping. While βS1 was not affected by R671C, homologous mutations in EmbS1 and PeriS1 significantly prolonged actin bound times to result in profoundly reduced detachment rates and altered load sensitivities.

### Homologous mutations had opposite effects on step size of single myosin molecules

Further analysis of HFS optical trapping data yields the step size of each myosin molecule in addition to its actin detachment kinetics. This analysis adapts the ensemble averaging method^55^ to HFS’s oscillatory data^44^. For each molecule, position traces of all events are aligned at the start of binding, extended to the length of the longest event, and averaged (Figure 5a). Myosin’s stroke is revealed after removal of the oscillations by subtracting a fitted sine function (Figure 5ab).

**Figure 5.**
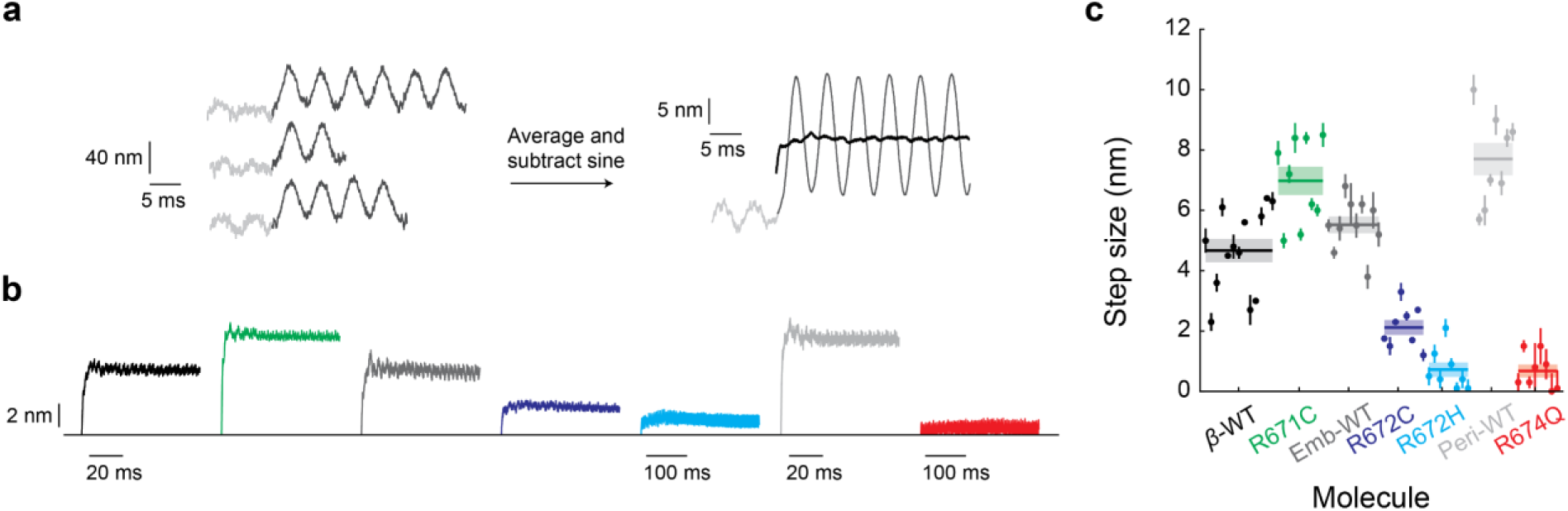
Single-molecule step sizes of myosins measured by optical trapping. **a.** Determination of step size from HFS data. Left: time traces of three example events before (light gray) and during (dark gray) binding. The amplitude of actin dumbbell oscillations increases upon binding to the myosin which is attached to the oscillating stage (see Supplementary Figure 5 and *Methods* for details of HFS). Right: hundreds of traces from one molecule are start-aligned, extended in time, and averaged (light and dark gray traces). A fitted sinusoid is then subtracted from the averaged trace to reveal the change in actin position due to myosin binding alone (black). **b.** Traces of step size from one molecule of each protein studied. **c.** Step sizes of all molecules measured. Each data point represents one molecule’s step size whose error bars represent uncertainties in the determination of change in actin position. Horizontal lines and shaded rectangles represent mean ± s.e.m. whose values are given in Table 1. All relevant pairwise comparisons are statistically significant except between βS1 WT and EmbS1 WT (Supplementary Table 1).

βS1 had step size *d* = 4.7 ± 0.4 nm, consistent with previous measurements in human β-cardiac myosin^44, 52, 56^. While EmbS1’s step size *d* = 5.5 ± 0.3 nm was not significantly different from βS1 (*p* = 0.088), that of PeriS1 was significantly larger (*d* = 7.7. ± 0.5 nm) (Figure 5ab, Table 1).

R671C increased the step size of βS1 by 49% (*d* = 7.0 ± 0.5 nm). On the contrary, the mutations in the developmental myosins greatly decreased myosin’s step size. R672C (*d* = 2.1 ± 0.3 nm) and R672H (*d* = 0.7 ± 0.3 nm) decreased the step size of EmbS1 by 62% and 87%, respectively, while R674Q all but eliminated PeriS1’s stroke (*d* = 0.7 ± 0.2 nm) (Figure 5ab, Table 1).

### Actin-detachment rate and step size of single myosin molecules account for effects of homologous mutations on ensemble motility

We next investigated the relationship between the single molecule optical trap and ensemble motility results. Conditions of our motility assay ensured that myosin heads were present in sufficient numbers on the surface and had relatively short tethers such that actin filaments’ sliding velocities were predominantly limited by their detachment from myosin rather than attachment^57, 58^ (Methods). Under these detachment-limited conditions, velocity can be approximated as the step size divided by the actin-bound time, which is inversely proportional to the detachment rate: *V* = *d*/*t*_b_ = *d*\**k*_det_. Indeed, velocities were linearly proportional to detachment rates (*R*^2^ = 0.81) (Figure 6a), confirming single molecule detachment kinetics as the basis of ensemble motility velocity. In contrast, linear regression of velocity versus attachment rate *k*_attach_ (see next section for calculation of *k*_attach_) gave *R*^2^ = 0.34 (Supplementary Figure S7).

**Figure 6.**
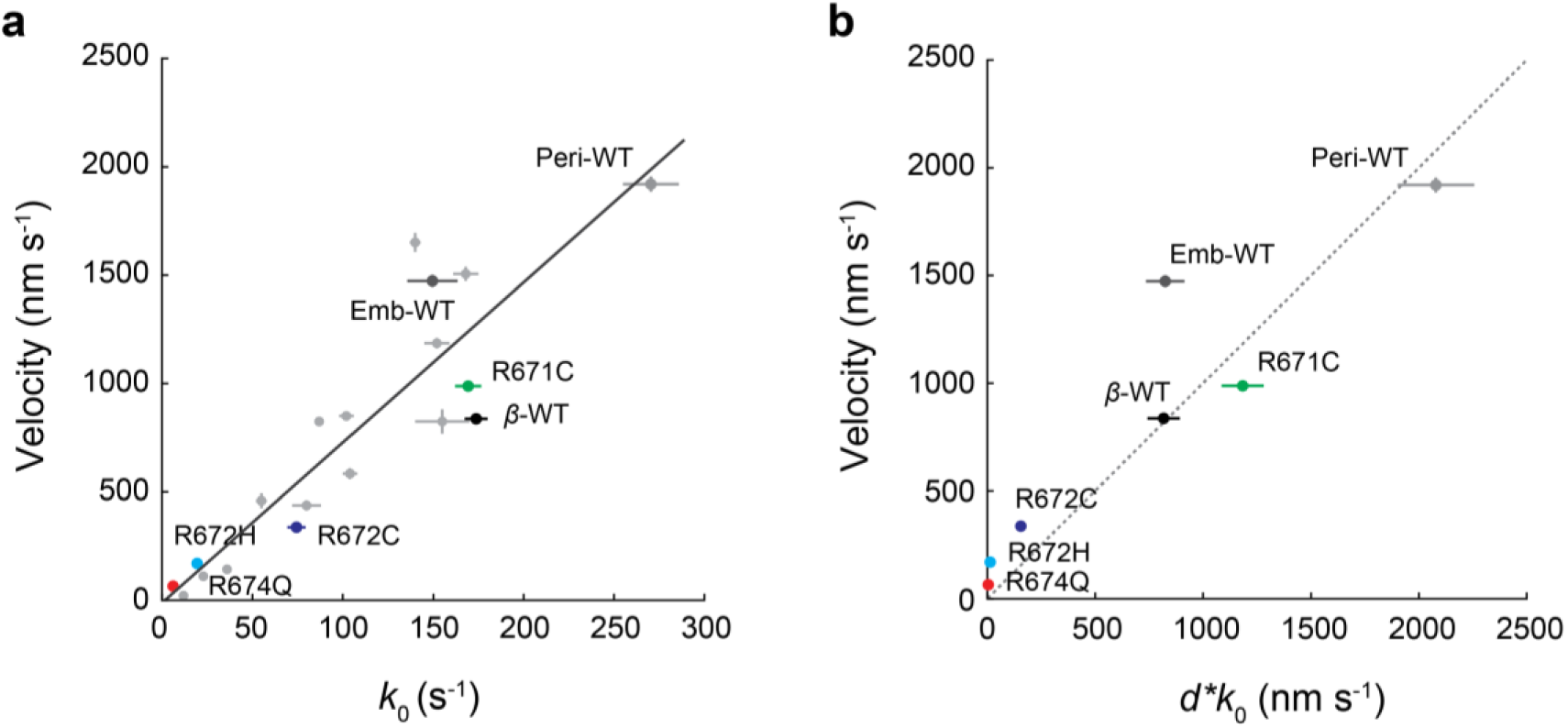
Single molecule detachment kinetics and step size determine ensemble actin sliding velocity. **a.** Velocities measured in the unloaded motility assay plotted against the detachment rate at zero force *k*_0_ at 23 °C (Table 1). Unlabeled gray data points are from Liu et al. *NSMB* 2018^42^. Error bars represent s.e.m. As velocity is predominantly detachment limited in our motility assay, linear regression gives a slope of 7.4 nm and *R*^2^ = 0.81. **b.** Measured velocities *V* (as in **a**) plotted against the product of measured step sizes *d* and measured detachment rates at zero force *k*_0_ (Table 1). Data from Liu et al. *NSMB* 2018 is not shown in **b** because step size data was not available. Horizontal error bars represent propagated errors in calculating *d*\**k*_0_. The dashed line represents *V* = *d*\**k*_0_.

As stipulated by the simple model *V* = *d*\**k*_det_, differences in the measured detachment rates and step sizes in general accounted for differences in the measured motility velocities (Figure 6b). The three constructs with the slowest velocities (R674Q PeriS1, R672H EmbS1, and R672C EmbS1) also had the slowest detachment rates and the smallest step sizes, in the same order. WT PeriS1 had the fastest velocity, consistent with it having the fastest detachment rate and largest step size. As another example, while R671C did not change βS1’s detachment kinetics, it did increase the step size, thus explaining the observed increase in velocity (Figures 3-6).

While single molecule detachment kinetics and step size were shown above as the basis for determining ensemble sliding velocity, we note that measured velocities in the unloaded motility assay were not exactly *d*\**k*_0_ (Figure 6b), as expected given factors involving both model oversimplification and experimental uncertainties. Motility velocities trended higher than *d*\**k*_0_, as also reported previously^42, 59^. EmbS1 WT (*V* = 1473 ± 28 nm/s, mean ± s.e.m.; *d*\**k*_0_= 825 ± 89 nm/s, propagated error), R672C (*V* = 337 ± 5 nm/s, *d*\**k*_0_= 155 ± 25 nm/s), R672H (*V* = 170 ± 2 nm/s, *d*\**k*_0_= 13 ± 6 nm/s), and PeriS1 R674Q (*V* = 65 ± 3 nm/s, *d*\**k*_0_= 4.3 ± 1.3 nm/s) serve as examples. Although the unloaded motility assay does not have externally added load like utrophin as used in loaded motility assays^38^, myosin molecules always experience load when working in an ensemble. Through a cycle of attachment, stroke, staying bound after the stroke, and detachment, a myosin head exerts both assistive and resistive forces on other heads bound to a shared actin filament. Myosin’s nonzero, asymmetrical load dependence δ (Figure 4) mechanically couples heads in an ensemble so that they detach and slide actin faster than single heads in isolation^59^.

We note R671C βS1 and WT PeriS1 as two exceptions to the trend above as their measured velocities were slightly lower than *d*\**k*_0_ (Figure 6b). Incidentally, these constructs required myosin concentrations higher than all other constructs in the motility assay to achieve smooth filament sliding (see Methods). It is possible that despite our efforts to provide conditions for detachment-limited motility, the combination of fast detachment kinetics, lower actin affinity (Table 1), and/or still-insufficient number of heads created an attachment-limited contribution that reduced velocity below *d*\**k*_0_. Furthermore, different groups of perturbations, such as hypertrophic vs dilated cardiomyopathy mutations or myosin activators vs inhibitors, may cause increased or decreased coordinated movement in ensemble than predicted by single molecule kinetics^42^.

### Duty ratio, average force and power predictions

Duty ratio is the fraction of time that myosin spends in the strongly bound, force-producing state and can be determined given the overall cycle rate *k*_cat_ measured by ATPase and the actin detachment rate measured by optical trapping. Since the detachment rate depends on load force, duty ratio is also function of load force *F*:

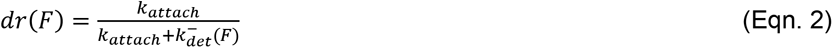

where *k_det_*^−^ (*F*)is the detachment rate calculated without harmonic force correction since we generalize to the non-oscillatory case:

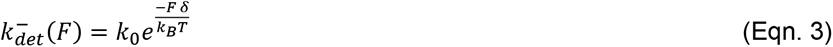

and the attachment rate *k*_attach_ was calculated at saturating actin concentrations and assumed to be independent of force:

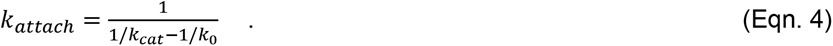

We highlight several features of the calculated duty ratios (Figure 7a, Table 1). First, duty ratio increased as load force increased due to the nonzero load dependence of the detachment rate: myosin remains bound to actin longer under increasing resistive loads. For most of the constructs measured, duty ratio was predicted to more than double as load is increased from 0 to 6 pN. The duty ratio of R672H EmbS1, on the contrary, did not increase to such extent since it was significantly less sensitive to load (Figure 4i, Table 1). Second, as expected of low duty ratio motors, most of these cardiac and skeletal muscle myosins had duty ratios below 5%, even under high resistive loads. Two exceptions were R672H EmbS1 and R674Q PeriS1 whose duty ratios under zero load (*dr*_0_) were 10% and 20%, respectively. This was because their detachment rates were much slower compared to other constructs (Figure 4h) while their overall cycle rate *k*_cat_ (Figure 2e) were not proportionally lower. Third, we note the effects of mutations. R671C did not change the duty ratio of βS1 (WT *dr*_0_ = 0.012 ± 0.002, R671C *dr*_0_ = 0.014 ± 0.003, propagated errors on calculated values) as it did not affect *k*_cat_ or *k*_det_. R672C and R672H increased the duty ratio of EmbS1 by 40% and ∼3.5x, respectively (WT *dr*_0_ = 0.028 ± 0.003, R672C *dr*_0_ = 0.039 ± 0.004, R672H *dr*_0_ = 0.102 ± 0.012). R674Q increased the duty ratio of PeriS1 10-fold (WT *dr*_0_ = 0.019 ± 0.007, R672C *dr*_0_ = 0.195 ± 0.043) owing to its dramatically prolonged bound time.

**Figure 7.**
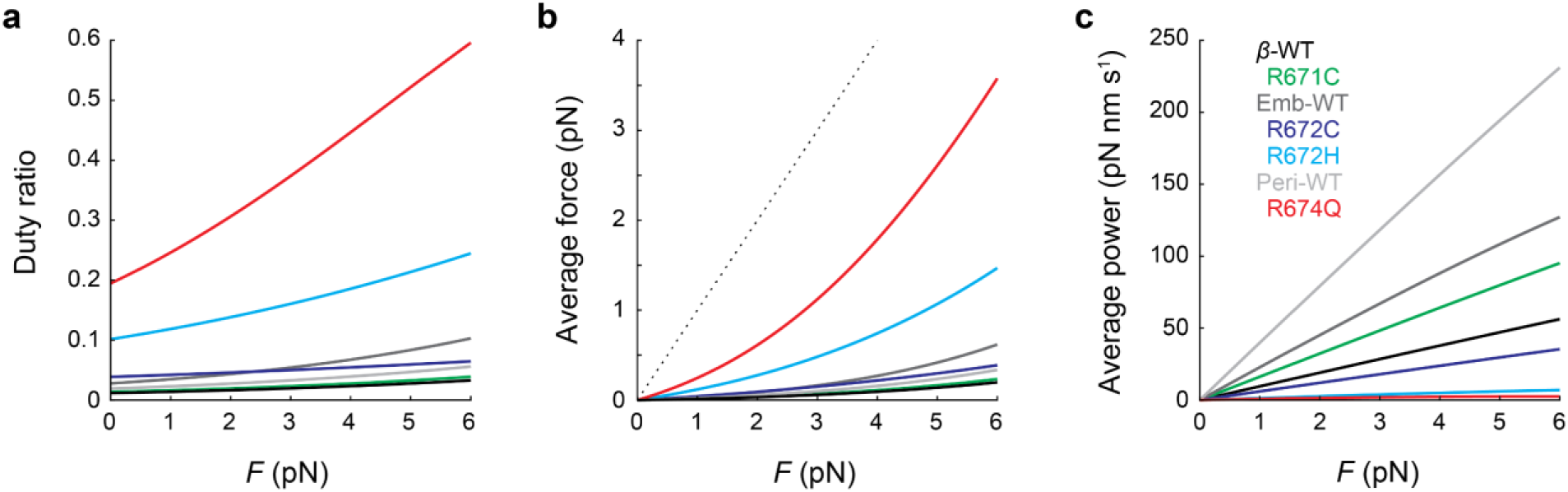
Duty ratio, average force, and average power of myosins determined by ensemble ATPase and single molecule measurements. **a–c.** Duty ratio **(a)**, average force **(b)**, and average power **(c)** as a function of the load force *F*, calculated by Equations (2)–(6). All curves are calculated at 30 °C using values of detachment rate kinetics (*k*_0_, δ), step size *d*, *k*_cat_, and *Q*_10_ given in Table 1. The values of duty ratio at zero load are also given in Table 1. Dotted line in **(b)** represents the average force if duty ratio = 1, in which case the average force equals the load force.

The average force produced by a single myosin molecule is equal to and opposite the load force *F* averaged over one cycle:

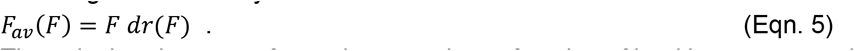

The calculated average forces increased as a function of load because myosin spends more time in the bound, force-producing state (Figure 7b). Myosins with higher duty ratios (eg. R672H EmbS1 and R674Q PeriS1) were predicted to produce higher average forces.

Finally the average power produced by a single myosin molecule is given by

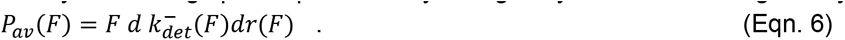

Because step size *d* is a determinant of power but not duty ratio, trends of the calculated average powers diverge from the trends of duty ratios and average forces (Figure 7c). WT perinatal and embryonic myosins had the highest powers due to a combination of large step sizes and fast kinetics. Despite having the same duty ratio, R671C was predicted to have higher power than WT βS1 due to the former’s larger step size. Finally, all three mutants in the developmental myosins were predicted to cause significantly decreased average power due to a combination of much reduced step sizes and slow kinetics.

### Simulations reveal a structural mechanism for step size and detachment rate reductions in embryonic R672C

How does one mutation at a highly conserved homologous site in closely related isoforms lead to such divergent molecular effects on motor function as revealed by our experimental results? As the answer must lie in myosin’s highly allosteric nature, we next explored this mechanism by using computational modeling and simulations to compare in particular R671C βS1 and R672C EmbS1, the same mutation in the two isoforms.

Given that the R672C mutation caused a reduction in step size and detachment rate in EmbS1 but not in βS1, we reasoned that this mutation must have a different impact on the structural or dynamic properties of the myosin motor depending on its broader sequence context. In most cases, experimental structures of point mutants reveal highly similar structures. On the other hand, a growing body of work supports the view that sequence variation can modulate the probability of adopting different conformations primed for specific functional roles, even when ground state structures are nearly identical^60–63^. When we constructed homology models of R672C EmbS1 and R671C βS1, we found very minor differences between structures. We reasoned, however, that R672C EmbS1 and R671C βS1 would adopt a different set of excited states that could explain their differences in biochemical and biophysical properties. Specifically, we hypothesized that R672C EmbS1 would reduce lever arm priming, reducing its ability to take a productive force-generating step on the actin track.

Hence, we performed molecular dynamics (MD) simulations of R672C EmbS1, WT EmbS1, R671C βS1, and WT βS1 motor domains starting from homology models of a fully primed pre-powerstroke crystal structure (PDB: 5N69 and 5N6A). In addition, we also performed simulations of WT βS1 starting from a homology model of a post-rigor crystal structure to compare the role of the arginine residue between different mechanochemical states. In total, we gathered ∼70 microseconds of data across 90 trajectories. To identify differences in the conformational ensembles of the 4 motors in their pre-powerstroke state, we utilized a deep learning method known as DiffNets^64^. DiffNets are self-supervised autoencoders capable of identifying features which distinguish ensembles in non-linear ways. Based on insight from a DiffNet (Supplementary Figure S8), we then measured the angle formed by the lever arm of our constructs relative to an experimental post-rigor structure (see Methods). We also assessed the opening of the nucleotide-binding site as well as the closing of the actin-binding cleft.

We find that the R672C mutation substantially reduces lever arm priming and ADP pocket opening in MD simulations of the Emb motor domain but that the same mutation is better tolerated in simulations of the β motor domain. We first investigated how the role of the highly conserved arginine residue changes between the post-rigor and pre-powerstroke states. R671 forms a network of hydrogen bonds in the pre-powerstroke state simulations that are significantly more likely than those same hydrogen bonds in the post-rigor state (Figure 8a). This suggests that R671 stabilizes the pre-powerstroke state and that loss of the arginine residue’s hydrogen bonds may destabilize this pre-powerstroke state. Indeed, we observed that mutations to cysteine at this position decrease the angle formed by the lever arm of our constructs relative to an experimental post-rigor structure in both EmbS1 and βS1 (Figure 8b, Supplementary Figure S9). However, EmbS1 and βS1 exhibit different behavior at the actin-binding cleft (Figure 8c, Supplementary Figure S10). In simulations started from a pre-powerstroke state model where the cleft is initially open, the cleft is more likely to close in R671 βS1 simulations than in R672C EmbS1 (Supplementary Figure S11). Because cleft closure is necessary for actin binding and a productive power stroke, we compared the position of the lever arm given a closed cleft. We find that among cleft-closed states, while wild type EmbS1 and βS1 occupy almost entirely lever-up states (∼85°), R672C EmbS1 occupies the lever-down (∼50°) states in a majority of simulations (Figure 8d). This significant loss in lever arm priming likely results in a less productive lever arm swing consistent with R672C’s experimentally measured large reduction in step size (Figure 5). In contrast, while R671C βS1 explores lever-down states with marginally higher propensity than wild type, it also exhibits a significant shift to higher-angle lever-up states (Figure 8d), again consistent with R671C’s experimentally measured increase in step size (Figure 5).

**Figure 8.**
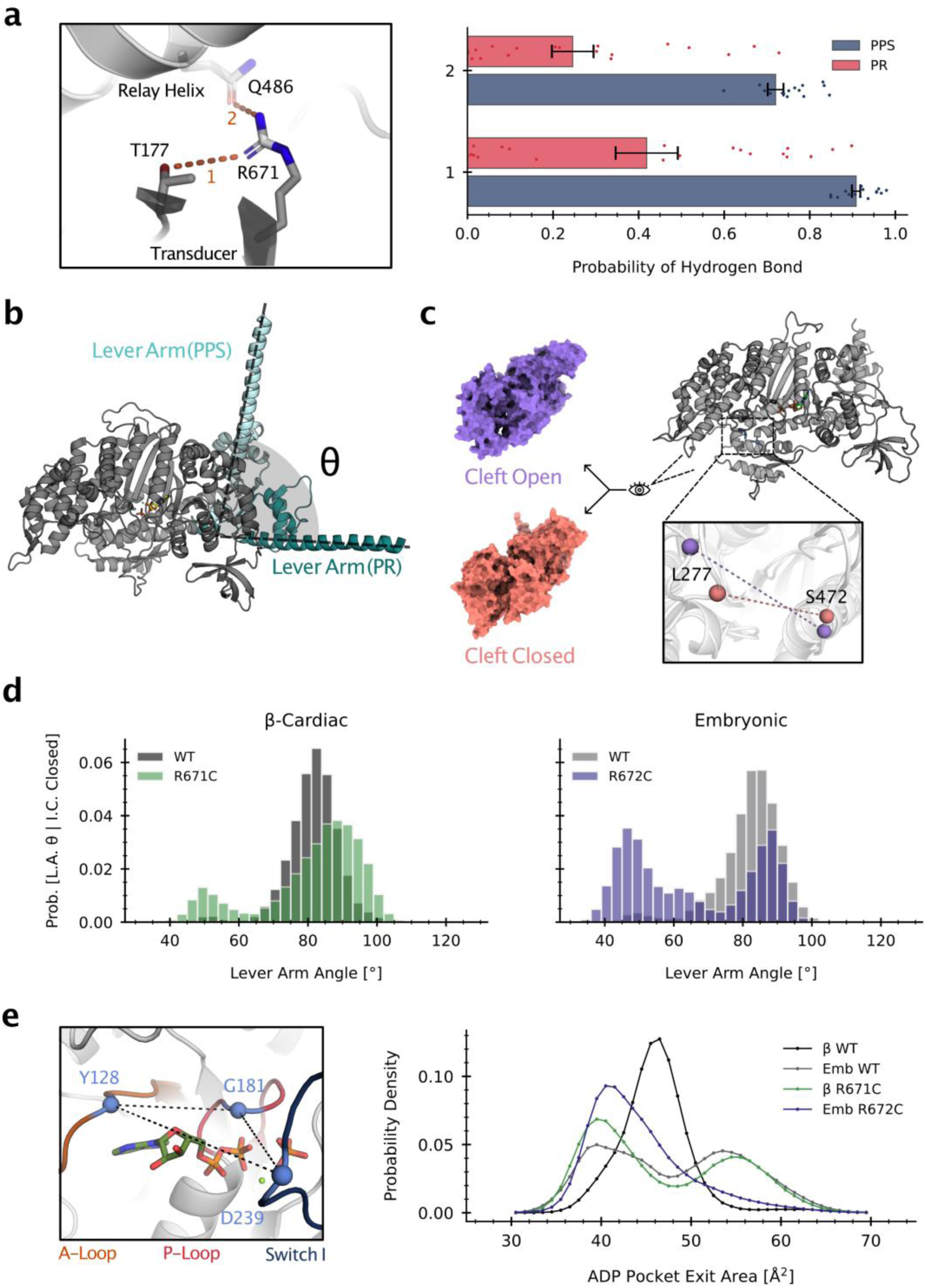
The R672C mutation reduces lever arm priming and ADP pocket opening in molecular dynamics simulations of embryonic myosin. All simulations were started from homology models built from template crystal structures of a pre-powerstroke wild type β-cardiac myosin. **a.** R671’s hydrogen bonds selectively stabilize the pre-powerstroke state in β myosin. A representative structure of the two hydrogen bonds is shown on the left. The bar chart (right) shows the probability of R671 forming hydrogen bonds with nearby residues T177 (transducer) and Q486 (relay helix) in simulations of the pre-powerstroke and post-rigor states. Data points represent individual trajectories and error bars are standard errors of the mean obtained from bootstrapping. **b.** Schematic depicting the lever arm angle that was measured in structures from simulations. A reference plane was defined by crystal structures of the pre-powerstroke (PDB: 5N69) and post-rigor (PDB: 6FSA) states. **c.** Opening and closure of the actin-binding cleft is observed in simulations. The inner cleft distance^94^ between the α-carbon atoms L277 and S472 (MYH7 numbering) is used as a metric of cleft closure. **d.** Among cleft closed states, Emb R672C exhibits a population shift towards lever-down states compared to Emb WT. Histograms show the distribution of lever arm angles conditioned on the binding cleft being closed (inner cleft distance ≤ 11 Å, Supplementary Figure S13). **e.** Emb R672C occupies ADP pocket-closed states more often than Emb WT. The ADP pocket exit area is defined by the area of the triangle formed by the α-carbon atoms of the three residues in the pocket (left). Simulations sample both pocket- open and pocket-closed states indicated by two peaks in the histogram (right).

Furthermore, we find that R672C EmbS1 predominantly occupies conformational states where the nucleotide-binding site is closed. We measured the opening of the ADP pocket by defining a triangle between the A-loop, P-loop, and Switch I (Figure 8e). While WT EmbS1 has a bimodal distribution of open and closed states (favoring closed), the R672C mutant shows a significant increase in probability density over closed states and a corresponding decrease over open states (Figure 8e, Supplementary Figure S12). As the release of ADP is a prerequisite for ATP binding and subsequent actin detachment, this finding may explain R672C EmbS1’s much reduced actin detachment rate observed in our single molecule trapping experiments (Figure 4h).

In summary, simulations reveal that the R672C mutation in EmbS1 selectively causes shifts in the distribution of pre-powerstroke conformations that likely contribute to the experimentally measured reductions in step size and detachment rate.

## Discussion

The R671C mutation’s lack of effect on several measures of βS1 motor function (ATPase (Figure 1a), actin detachment rate and force sensitivity (Figure 4d)) is reminiscent of many (R453C^37^; R403Q^65^; the converter domain mutations R719W, R723G, and G741R^66^; V763M^32^; R663H^67^), but not all (the early-onset mutations H251N and D239N^41^), previously studied HCM mutations (reviewed in Kawana *Front Physiol* 2022^33^). However, R671C is to date unique in that it increased step size (Figure 5), whereas previously only unchanged (R453C^37^, R403Q^65^, D778V and S782N^46^) or decreased (P710R^44^, R712L^68^, L781P^46^) step sizes have been reported. Simulations suggest that the larger step size may be the result of a further primed lever arm in the pre-powerstroke state (Figure 8d). This larger step size results in a higher predicted power output despite an unchanged duty ratio (Figure 7). Beyond the motor domain, future studies employing an S2-containing construct which enables formation of an autoinhibited state^69, 70^ may find that R671C reduces autoinhibition, which, in combination with its increased step size, would form the basis of molecular hypercontractility for this HCM mutation.

In contrast to R671C in βS1, the homologous mutations in the developmental skeletal myosins had much greater effects on all measures of motor function: ATPase (Figure 1bc), actin detachment rate and its force sensitivity (Figure 4ef), and step size (Figure 5). The prolonged actin bound times and reduced step sizes of the embryonic and perinatal mutants are reminiscent, to different degrees, of the investigational heart failure drug omecamtiv mecarbil (OM)^71^. OM reduced the actin detachment rate of β-cardiac myosin by ∼10 fold (from ∼100 s^-1^ to ∼10 s^-1^)^42, 52^ and demolished its step size^52^ (Spudich lab unpublished data). Despite these inhibitory effects on the myosin molecule, OM activates the heart muscle through a proposed mechanism in which the prolonged actin bound times enable myosin heads to cooperatively keep the regulated thin filament open at lower calcium levels, thus prolonging contraction^42, 52^. Indeed, our lab’s unpublished results show that OM increases the calcium sensitivity of regulated thin filaments in the motility assay. Future studies may reveal whether the developmental skeletal myosin mutants also have increased calcium sensitivity at the molecular level due to their prolonged actin bound times.

Comparing myosin motor dynamics provides insight into how the same mutation can have different, or even opposite, effects depending on its sequence context. Predictions based on homology modeling alone^15^ would suggest that homologous mutations would have the same effect on motor function. For example, a previous study using homology modeling had suggested that the R671 residue is at a communication hotspot between the nucleotide-binding site and the lever arm, so mutations that perturb this communication would disrupt aspects of force production^28^. While our findings of much reduced force sensitivity, actin detachment rate, and step size in R672C EmbS1 provide experimental evidence for this prediction, the same mutation in βS1 (R671C) has a larger step size and no effect on the load-dependent actin detachment kinetics. In molecular dynamics simulations launched from the same starting structure, R672C EmbS1 is more likely to adopt structures with a closed nucleotide-binding site than R671C βS1. R672C EmbS1 is also more likely to adopt structures with the lever arm in a less primed position when its actin-binding cleft is closed. Thus, MD simulations of each mutant in its specific sequence background are necessary to uncover differences in the distribution of conformations adopted by myosin motors, in order to explain functional differences among homologous mutations^61–63^. Future simulation studies will address other homologous mutations for which experimental data exist and incorporate actin, regulated thin filament proteins, etc. to the models to directly probe how myosin behaves when bound to actin.

Our findings of the effects of the mutations at the molecular level provide the mechanisms underlying prior observations in muscle cells. A previous study had found that FSS patient-derived muscle cells and myofibrils containing embryonic myosin bearing the R672C mutation have ∼3- 5x slower rates of relaxation than controls ^24^. Our results provide direct evidence of slowed actin detachment kinetics at the single molecule level as the basis for slowed relaxation in cells. MD simulations further provide insight that a smaller ADP exit area may contribute to the longer actin-bound times observed in R672C EmbS1. In another study, FSS mutations (Y583S and T178I) in *Drosophila* skinned muscles increased stiffness by 70-77% and decreased power by 50-66%, and the authors proposed longer bound times as the underlying mechanism^25^. Future optical trapping measurements of the load-dependent detachment kinetics of these mutants will determine whether that is indeed the case.

More importantly, our results reveal a partial, nuanced correlation between the magnitude of changes due to mutations and disease severity. Within the embryonic isoform, the correlation is clear. The histidine mutation causes a more severe disease than cysteine in FSS patients^23^, a trend mirrored by our molecular findings: R672H had larger reductions in ATPase *k*_cat_, actin- detachment rate, and step size than R672C. When comparing across the isoforms, the trend is more nuanced. First, FSS is considered to be the most severe of the distal arthrogryposis syndromes, more so than TPS, but the TPS R674Q mutation in perinatal myosin had the largest effects out of all mutations. Second, in one view, HCM may be considered the most serious of all three myosinopathies because it increases the risk of atrial fibrillation, heart failure, and sudden cardiac death; indeed, it is the leading cause of sudden cardiac death in young athletes^72^. But in a different view, while most patients with FSS and TPS have a normal life expectancy, their clinical signs are arguably much more visually striking than those with HCM. Thus, the HCM R671C mutation’s relatively small effect (enhanced step size) in β-cardiac myosin correlates well with an HCM heart’s relatively mild phenotype. A mutation in β-cardiac myosin that drastically changes its motor function would be catastrophic in the heart and thus not observed in the population.

Understanding the effects of different mutations on various parameters of motor function enables development of small molecule therapeutics that specifically reverse those changes at the source rather than drugs that act downstream. Prolonged actin-bound times may be a common defect of developmental skeletal myosins implicated in FSS, TPS, and other congenital contractures. In such cases, a logical candidate is 2-deoxy-ATP as it has been shown to increase the actin detachment rate of single β-cardiac myosin molecules by ∼70%^42^ and improve muscle contractility^73^. There are other small molecule myosin effectors that have been shown to prolong actin bound times, alter step size, change the overall ATP cycle rate, inhibit actin binding, and/or promote the super-relaxed state^42, 52, 74–76^.

While the developmental muscle myosins cease to be expressed in significant levels in almost all skeletal muscles shortly after birth, their devastating effects clearly outlive their transient expression. The significance of longer actin bound times, altered force sensitivities, and reduced step sizes in the embryonic and perinatal mutants may be appreciated in the context of the critical role of mechanosensation in fetal muscle development^18^. The shared defects of dramatically longer bound times and reduced step sizes likely play major roles in disease mechanisms. As both lower (R672C and R672H EmbS1) and higher (R674Q PeriS1) force sensitivities lead to similar diseases involving muscle contractures, the precise mechanism by which altered myosin force sensitivity disrupts development remains to be elucidated in future studies.

In the large myosin superfamily of motor proteins, each isoform has distinct mechanical and kinetic properties tuned to its specific function, as transporters, force sensors, or drivers of muscle contraction. Among muscle myosins, the motor properties of the isoform determine the identity of the particular muscle type (eg. cardiac, skeletal, smooth). Despite these differences, myosins share a common global structure and perform the same function of actin-based motility through ATP hydrolysis. Perturbations in key conserved regions of the molecule are then expected to be disruptive across isoforms, as is the case for mutations at the homologous R671 residue in β, embryonic, and perinatal myosins. By measuring key factors of molecular power generation, we have found that while each mutation had distinct effects, they affected the developmental myosins much more than β myosin. Furthermore, the observed molecular changes correlate partially with clinical disease severity. Molecular dynamics simulations provide possible structural explanations for the differentially affected properties. Our findings of divergent effects on motor function due to homologous mutations present a striking illustration of myosin’s highly allosteric nature.

## Materials and Methods

### Expression and purification of proteins

Recombinant human myosin S1 constructs for MYH7(Met^1^–Ser^842^), MYH3(Met^1^–Ala^844^), MYH8(Met^1^–Leu^842^) containing a GSG-RGSIDTWV C-terminal tag were cloned into a pUC19 vector, and site directed mutagenesis was performed to generate the 4 mutations described herein. The transgenes were inserted into pShuttle-CMV of the AdEasy Adenoviral Vector System (Agilent #240009) which was used to produce recombinant adenovirus of the seven constructs. A recombinant adenovirus of the human essential light chain (ELC or MYL3) with a 6×His N-terminal tag^31^ was also used. Adenovirus was amplified in HEK293 cells and purified from clarified cell lysates in a CsCl step gradient followed by a linear gradient. The concentration of recombinant adenovirus was determined from OD_260_ measurement of a 1:100 dilution and conversion factor of 1 OD_260_ = 1.1 × 10^12^ viral particles/ml. Virus stocks were stored at −20°C in 80 mM Tris pH 7.5, 200 mM NaCl, 0.8 mM MgCl_2_, 0.8 mg/ml BSA, 40% glycerol.

Murine C_2_C_12_ myoblasts were cultured in Dulbecco’s modified Eagle’s Medium (DMEM) supplemented with 100 units/mL Penicillin, 10 ug/mL Streptomycin, 2 mM L-glutamine, and 10% fetal bovine serum (FBS) in four-layer Nunc^TM^ Cell Factory systems, and incubated for 3 days at 37°C, 5% CO_2_ until 100% confluent. Cells were differentiated into myotubes by changing media to 2% Horse Serum in place of FBS, followed by incubation for 3 days at 37°C, 5% CO_2_. Myotubes were co-infected with 2 × 10^9^ viral particles/mL of each virus (heavy chain + MYL3 light chain) diluted in media containing 5% FBS, and incubated for 5 days at 37°C, 5% CO_2_. Cells were detached by treatment with trypsin and collected in 120 mL per factory of 30mM HEPES, 135mM NaCl, 15mM KCl, 2mM EDTA, 60mM Sucrose, 0.2% Pluronic F-68, 0.5 mM AEBSF, pH 7.5. Pellets were frozen in LN_2_ and stored at -80°C.

Myosin was purified through two rounds of affinity chromatography, first using an affinity clamp (PDZ)^77^ coupled to Sulfolink™ Coupling Resin (Thermo #20401) to capture the myosin S1 heavy chain by the 8 amino-acid tag (“affinity clamp”, AC) (S1-AC) at its C-terminus, followed by nickel resin to capture the recombinant human ELC by its 6×His tag, allowing myosin constructs bound to native mouse ELC to be removed. Cell pellets were resuspended in 8 mL lysis buffer (50mM Tris, 0.2M NaCl, 4 mM MgCl2, pH 8.0, 2 mM DTT, 1 mM ATP, 0.5% (v/v) Tween, 0.2 mM phenylmethanesulfonyl fluoride (PMSF), 1X Roche cOmplete EDTA free tablet) per gram pellet and lysed by dounce homogenization 15-20X on ice. Lysate was clarified at 40,000 x g for 25 min at 4°C, and supernatant was passed through a 5 uM then 1.2 uM filter before batch binding to PDZ-coupled resin with agitation for 30 minutes at 4°C. Resin was packed into a chromatography column and 10 column-volumes (CV) of wash buffer (20 mM HEPES, pH7.5, 50 mM KCl, 5 mM MgCl_2_) with 5 mM DTT and 1 mM ATP were flowed through. Myosin was eluted from the resin in 10 CV of wash buffer containing 0.1 mM DTT and 0.1 mg/mL of an elution peptide (WETWV). This was collected and passed over Nickel resin pre-equilibrated in wash buffer. Myosin with human ELC bound was eluted from nickel resin in wash buffer with 400 mM imidazole, and fractions containing myosin were pooled and dialyzed overnight against wash buffer with 10% sucrose and 5 mM DTT. Myosin was removed from dialysis the next day, ATP was added to 1 mM, and myosin was frozen in aliquots in LN_2_ and stored at -80°C.

### Actin-activated ATPase

ATPase activity was measured by an NADH-coupled assay^78^. G-actin was polymerized to F-actin the day before the assay by dilution to 250 uM with assay buffer (20 mM HEPES, 50 mM KCl, 5 mM MgCl2, pH7.5). The next day gelsolin was added to 1 uM and incubated with F-actin for 30 min before titrating with assay buffer. Actin titrations were added to a 96-well plate and mixed with myosin, followed by incubation for 10 minutes before the reaction was initiated by addition of a 10X Reagents solution. The plate was shaken for 1 minute before reading the absorbance at 340 nm every 30 sec for 1 hour. Reactions were carried out at 30°C in a 200 uL volume containing 0- 100 uM actin, 0.2 uM myosin, 20 mM HEPES, 50 mM KCl, 5 mM MgCl2, pH7.5, 5 mM DTT, 3 mM ATP, 3 mM PEP, 1 mM NADH, and 3-7 units/mL of pyruvate kinase and lactate dehydrogenase (Sigma P0294). An NADH standard curve was measured in order to convert absorbance values to the concentration of ATP hydrolyzed. Mutant myosin was run on the same plate as its WT counterpart in technical triplicate for each actin concentration measured. ATPase rates were determined from the slope of the absorbance (ATP hydrolyzed) per time for each well, divided by the concentration of myosin to obtain a /sec rate, which was then plotted against actin concentration. The data was fitted to the Michaelis-Menten equation for enzyme kinetics to determine k_cat_ and K_m_ for each biological replicate myosin. Average values and SEM of k_cat_ and K_m_ from 3 biological replicates are reported in Table 1, and t-tests were used to determine significance between measured parameters of myosin pairs in Supplementary Table 1.

### Deadheading myosin

Myosin protein used for unloaded motility and optical trap experiments was further subjected to a ‘deadheading’ procedure to remove inactive heads, as commonly performed and reported in the literature. Myosin at 1uM was mixed with 3 uM unlabeled F-actin on ice for 5 min, followed by addition of 3 mM ATP, then centrifuged at 95 krpm in a TLA-100 rotor (Beckman Coulter) at 4°C for 20 min. The supernatant was collected. We found that the β R671C protein had more inactive heads than β WT, as seen in a greater percentage of stuck filaments (∼15% for R671C vs ∼7% for WT, Supplementary Figure S4). Embryonic WT and mutants had low percentages of stuck filaments (0-5%). Perinatal WT had a greater percentage of stuck filaments than R674Q (19% for WT vs 6% for R674Q). Note, however, that these stuck filament percentages do not directly translate to the percentage of inactive heads; only a very small number of inactive heads (probably 1% or less) is needed to produce a much larger number of stuck filaments. In addition, deadheading cannot remove 100% of completely or partially inactive heads.

### In vitro motility

Motility measurements used our previously described motility assay^38, 79^ with some modifications. Multi-channel flow chambers consisted of double-sided tapes between a glass slide and a coverslip pre-coated with a solution of 0.1% nitrocellulose/0.1% collodion in amyl acetate.

The following solutions were flowed into the chamber sequentially: 1. SNAP-PDZ18 (purified as described in Aksel et al. *Cell Reports* 2015) diluted to 3 μM in assay buffer (AB; 25 mM imidazole pH 7.5, 25 mM KCl, 4 mM MgCl_2_, 1 mM EGTA, and 10 mM DTT), incubated for 2 min. 2. BSA diluted to 1 mg/mL in AB (ABBSA), incubated for 2 min. 3. S1-AC diluted to ∼200 nM (except as indicated in the paragraph below) in ABBSA, flowed twice with 2 min incubation each time. 4. ABBSA. 5. A final solution consisting of 2 mM ATP, 2.5 nM TMR-phalloidin-labeled actin filaments, and an oxygen-scavenging system (0.4% glucose, 0.11 mg/mL glucose oxidase, and 0.018 mg/mL catalase) in ABBSA. pH of the final solution was measured to be 7.5 at 23 °C. See Movies S1-S14 for example motility measurements of each protein. These conditions ensured that enough heads were on the surface such that velocities were detachment-limited. This is supported by the relationship between filament length and velocity in Figures S2 and S3 and by the absence of “wavy” filaments (in search of myosin heads) in the Movies. The concentration of myosin we used was empirically optimized; using too high a concentration did not increase velocities further and resulted in excessive shredding of actin filaments into very short fragments.

Two exceptions to using ∼200 nM myosin were as follows. 1. Undiluted β R671C S1-AC was used, as dilution of the protein after purification resulted in wavy filaments in motility, indicating insufficient myosin heads on the surface. While we estimated the concentration to be ∼500 nM, accurate determination of protein concentration at low levels was difficult as this mutant expressed and purified relatively poorly. 2. 400-600 nM perinatal WT S1-AC was used to achieve non-wavy filament movement. Accurate determination of concentration was not an issue in this case. Instead, we attribute the slightly higher concentration requirement to perinatal WT’s lower affinity for actin (higher *K*_M_, Figure 2) and much faster actin-detachment rate (Figure 4F and H). Since we do not use a viscous agent like methylcellulose to artificially keep filaments confined to the surface, more myosin heads are likely necessary to keep filaments on the surface when the actin-bound times are very short.

The slide was then placed on a Nikon Ti-E inverted microscope with a 100x TIRF objective. Images were taken for 30-60 sec at 0.5-2 Hz with 200-300 ms exposure on an Andor iXon+EMCCD camera. Different imaging rates and durations were used depending on the actin sliding velocities to accurately capture frame-to-frame filament movements while minimizing file size. Three movies were recorded for each channel. The objective temperature was maintained at 23 +- 0.3 °C or 30 +- 0.3 °C as closely as possible to minimize variations in velocities stemming from myosin’s high sensitivity to temperature. This attention to precise temperature contributes to the low variance in velocities as apparent in Figure 3. Movies of all channels on each slide were acquired as quickly as possible, within 15 min after flowing in the final buffer. Despite our foregoing an ATP regeneration system, no slowing down of actin filaments was observed, nor any differences observed with or without an ATP regeneration system.

Movies were processed by FAST (Fast Automated Spud Trekker) for filament tracking and velocity analysis^38^ (Supplementary Figures S2 and S3). The following parameters were used in FAST: window size *n* = 5, path length *p* = 10, percent tolerance *pt* = 20, minimum velocity for stuck classification *minv* = 80 nm/s, overlap score *oscore* = 0.3, and log area score *lascore* = 1.3. Window size is the number of consecutive frames *n* over which frame-to-frame instantaneous velocities are averaged (let *v*_n_ be this n-frame-averaged velocity). FAST outputs each *v*_n_ rather than one velocity averaged over a filament’s entire path. Path length is the minimum number of frames in which a filament must be present in order for that filament’s *v*_n_’s to be included in the analysis. Percent tolerance *pt* is a parameter for further filtering out *v*_n_’s with non-smooth motion, as follows: *v*_n_’s are filtered out if their standard deviation (within the n frames) is greater than *pt*/100\**v*_n_. Minimum velocity for stuck classification *minv* is the minimum value (in nm/s) of the average of all *v*_n_’s for one filament for that filament to be classified as “stuck”, in which case all of the *v*_n_’s for that filament are then set to 0. Finally, *oscore* and *lascore* are two advanced parameters for accurate filament tracking whose default values (0.4 and 1, respectively) were typically used in previous publications. The overlap score *oscore* is the minimum dot product between 2 filaments of adjacent frames in order for those frames to be included. The log-area score *lascore* is the maximum cross product between 2 filaments of adjacent frames in order for those frames to be included. In experiments for our current paper using fast myosins at higher temperatures (eg. perinatal WT and embryonic WT at 30 °C), the higher velocities movies required slightly more relaxed values (0.3 and 1.3, respectively) for proper filament tracking; otherwise, many velocities were incorrectly filtered out as the frame-to-frame filament displacement was large when velocities were high. For consistency, we analyzed all movies using the same FAST parameters.

“Mean velocity including stuck filaments (MVIS)” is the mean of all *v*_n_’s (including those that are 0) that satisfy the path length criteria. MVIS is not affected by the chosen percent tolerance value. Note that MVIS is equivalent, up to a multiplicative factor, to the “percent time mobile” metric defined previously (9). “Mean velocity (MVEL)” is the mean of *v*_n_ > 0 and which satisfy the path length criteria. “Mean velocity – filtered (MVEL_pt_)” is the mean of *v*_n_ > 0, which satisfy the path length criteria, and which satisfy the tolerance criteria (call these *v*_n_’s “filtered velocities”). All velocities reported in the main text and figures are MVEL_20_. Note that as the value of *pt* increases, MVEL_pt_ approaches MVEL. “Top5%” is the mean of the highest 5% values of “filtered velocities.” “Stuck percentage (%STUCK)” is the percentage of *v*_n_’s that are equal to 0.

TOP5% or MVEL_20_ using *minv*=80 and *pt*=20 are appropriate for analysis of unloaded motility data, as reported in some previous publications. In other previous publications^44^, we have also reported MVIS using lower *minv* value and higher *pt* value as appropriate for analyzing loaded or regulated-thin-filament motility data in which filaments have stop-and-go movements. While we report MVEL_20_ in the main Figure 3 of this paper, we provide TOP5% MVIS, and %STUCK in Figure S4 for comparison. Relationships of velocities between the seven proteins are highly conserved regardless of the velocity metric used.

Statistical significance was determined using a two-tailed unequal variances t-test.

### Single molecule optical trap measurements

The load-dependent detachment rates of myosin S1-AC molecules were measured in a dual-beam optical trap using the harmonic force spectroscopy (HFS) method previously described^42, 45, 80^ with slight modifications. The sample chamber consisted of two double-sided tapes between a glass slide and a coverslip spin-coated with 1.6-um-diameter silica beads (Bang Laboratories) and a solution of 0.1% nitrocellulose/0.1% collodion in amyl acetate. Experiments were done at 23 °C. The following solutions were flowed into the chamber sequentially, each with a couple minutes of incubation: 1. SNAP-PDZ diluted to 5-20 nM in assay buffer (AB; 25 mM imidazole pH 7.5, 25 mM KCl, 4 mM MgCl_2_, 1 mM EGTA, and 10 mM DTT). 2. BSA diluted to 1 mg/mL in AB (ABBSA), to wash and block the surface. 3. Myosin S1-AC diluted to 50-300 nM in ABBSA, to bind all the SNAP-PDZ18 on the surface. 4. ABBSA, to wash. 5. A final solution consisting of 2 mM ATP, 0.3 nM TMR-phalloidin-labeled biotinylated actin (Cytoskeleton) filaments, an oxygen-scavenging system (0.4% glucose, 0.11 mg/mL glucose oxidase, and 0.018 mg/mL catalase), and 1-um-diameter NeutrAvidin-coated polystyrene beads (ThermoFisher) or 1-um-diameter streptavidin-coated polystyrene beads (Bang Laboratories) diluted in ABBSA. The chamber was sealed with vacuum grease and used for up to 2 hr. A range of SNAP-PDZ and myosin concentrations are given above because these concentrations were varied empirically to achieve clean binding events under single molecule conditions for different constructs.

The two traps were calibrated using the power spectrum method with corrections for filters, aliasing, and surface effects^81^, and their stiffnesses were 0.09-0.13 pN/nm in these experiments. After calibration, a “dumbbell” was made by stretching an actin filament between the two trapped beads (Figure 4A). While the stage oscillated at 200 Hz, the dumbbell was lowered near platform beads on the surface to test for binding to a potential myosin. Robust interactions with myosin were indicated by consistent, large, brief displacements of the trapped beads due to stronger association with the oscillating stage. We typically explored ∼4-10 platform beads before robust interactions were observed, suggesting that the SNAP-PDZ concentration used resulted in sufficiently low numbers of properly oriented molecules on the surface for single molecule conditions. Time traces were saved and automatically processed as previously described^42, 45^. Briefly, events were selected based on a simultaneous increase in oscillation amplitude and decrease in phase between each bead and the stage (Supplementary Figure S5). The detachment rate at each force was determined by a maximum likelihood estimation from the durations of events within the force bin. The rates at different forces were then fitted to Eqn. 1 to obtain the two parameters *k*_0_ and δ. All binding events and detachment rates of one example molecule for each of the seven proteins are shown in Supplementary Figure S6. The data on multiple molecules presented in this paper are from multiple dumbbells in independent experiments from multiple days. Statistical significance was determined using a two-tailed unequal variances t-test.

On a note to clarify any potential questions, 200 Hz stage oscillations set the minimum detectable events at 5 ms. PeriS1 had *k*_0_ ∼ 270 s^-1^, which corresponds to average bound times of ∼4 ms, and assistive forces result in even shorter events. However, we are able to measure these rates, with appropriately larger uncertainties as presented, because the bound times are exponentially distributed. While events shorter than 5 ms are missed, there is a sufficient number of events longer than 5 ms (Supplementary Figure S6) to determine rates from the maximum likelihood fits.

The step sizes of myosins were determined from the same HFS data by adapting the ensemble averaging method to HFS’s oscillatory data^44, 55^. For each molecule, position traces of all events were aligned at the start of binding, extended to the length of the longest event, and averaged (Figure 5A). Then the oscillations were removed by subtracting a fitted sine function (Figure 5A). The total step size for each molecule was taken as the difference between the initial position and the end position of the extended, averaged, sine-subtracted traces (Figure 5B). Error bars represent uncertainty in the determination of this difference in positions. Statistical significance of differences between proteins was determined using a two-tailed unequal variances t-test.

### MD simulations

All simulations were performed using GROMACS 2022.4^82^ on our in-house commodity cluster using a mixture of NVIDIA RTX A5000, Quadro RTX 6000, and Tesla P100 GPUs. The protein component and ADP of the motor domains were parameterized using CHARMM36^83^. Inorganic phosphate was parameterized using the Karplus parameters^84^. Magnesium was parameterized using the improved parameters developed by Allner et al.^85^. The protein structure was solvated in a dodecahedral box of TIP3P water^86^ that extended 1 nm beyond the protein in every dimension. Thereafter, sodium and chloride ions were added to the system to maintain charge neutrality and 0.1 M NaCl concentration. Each system was minimized using steepest descent until the maximum force on any atom decreased below 1000 kJ/(mol x nm). The system was then equilibrated with all heavy atoms restrained in place at 310 K maintained by the Berendsen thermostat^87^ and the Parrinello-Rahman barostat^88^. Production simulations were run in the NPT ensemble at 310 K using the leapfrog integrator, Bussi-Parinello thermostat^89^, and the Parrinello-Rahman barostat^88^.

Simulations of the pre-powerstroke state began from homology models built from crystal structure templates (PDB: 5N6A and 5N69) using MODELLER^90^. Simulation lengths varied from 500 ns-1.5 µs. The aggregate simulation time in the study was ∼70 µs. Simulations of the post-rigor state in wild-type β myosin were started from homology models created by MODELLER from a post-rigor crystal structure template (PDB: 6FSA)^91^.

Trajectory analysis was conducted in Python using the library MDTraj^92^. A DiffNet was trained using heavy atoms out to C-gamma for all 4 constructs using standard training parameters (EM bounds of [0.1,0.4] and [0.6,0.9], 150 latent variables, and binary initial labels)^64^. Data visualization was completed using matplotlib. Gaussian mixture models were fitted using the implementation of the EM algorithm in scikit-learn. Structures were visualized in PyMol or UCSF ChimeraX.

The lever arm angle for each frame was computed by aligning each simulation frame to a post-rigor crystal structure (PDB: 6FSA) and measuring the angle between a simulation structure’s lever arm and the post-rigor crystal structure in the plane formed by the post-rigor crystal structure and a pre-powerstroke crystal structure (PDB: 5N6A). Calculations were made in this plane to ensure that we only considered axial angular deviation rather than azimuthal deviation^93^. The lever arm angle was calculated by first determining a vector between the beginning (center of mass of C-alphas from residues 769-774, MYH3 numbering) and end (center of mass from C-alphas from residues 775-780) of our lever arm constructs. Then, an axial plane was defined based on a vector from the beginning of the post-rigor lever arm (center of mass (COM) of C-alphas from residues 769-776 from 6FSA) to the end of the post-rigor lever arm (C-alpha COM from 779-787 from 6FSA) and another vector from the beginning of the post-rigor lever arm to end of the pre-powerstroke lever arm (C-alpha COM from 766-772 from 5N6A). Finally, the angle between the simulation frame’s projected vector onto the axial plane and the post-rigor structure lever arm vector was calculated.

To determine a threshold for defining when the actin-binding cleft was closed, we determined cleft distances in actomyosin rigor structures and performed a sensitivity analysis. We find that the inner cleft distances for β myosin range from 0.94 nm to 1.0 nm (PDB: 7JH7, 8EFH, 8EFI). Given that there is some role for induced fit in actin binding^94^ and because we had very few simulation frames that achieved these small cleft distances, we decided on a threshold of 1.1 nm. We demonstrate that R672C EmbS1 is more likely to adopt lever arm down states than R671C βS1 across all inner cleft thresholds in a sensitivity analysis (Supplementary Figure S13).

## Acknowledgements

C.L., K.M.R., and J.A.S. were funded by NIH Grants R01 GM033289, RM1 GM131981, and HL117138 to J.A.S. C.L. was also funded by the postdoc career fund at Lawrence Livermore National Laboratory. A.K. and L.A.L were funded by NIH Grants 5T32 HL007822 and R01 GM29090 to L.A.L. A.M. was funded by NIH F30 Fellowship 1F30HL162431-01A1. G.R.B. was funded by NSF CAREER Award MCB-1552471, NIH Grants R01 GM124007 and RF1AG067194, and Packard Fellowship for Science and Engineering from The David & Lucile Packard Foundation. The microscope was funded by NIH Grant S10RR026775 to J.A.S.

## Author contributions

C.L. performed motility and trap experiments and analyzed data. A.K. performed molecular cloning, viral production, protein expression and purification, ATPase experiments, and analyzed data. A.M., A.B., and C.J.A. performed simulations and analyzed data. C.L., A.K., A.M., A.B., L.A.L., and K.M.R. wrote paper. All authors contributed intellectually to discussions of the results and reviewed and edited the paper.

## Supplementary Movies

### Examples of motility movies

Compiled movies are all ∼4x speed

1. beta WT 23C playback speed 3.7x 7fps
2. beta R671C 23C playback speed 3.7x 7fps
3. Emb WT 23C playback speed 3.7x 7fps
4. Emb R672C 23C playback speed 3.7x 7fps
5. Emb R672H 23C playback speed 4x 4fps
6. Peri WT 23C playback speed 3.7x 7fps
7. Peri R674Q 23C playback speed 4x 2fps
8. beta WT 30C playback speed 3.7x 7fps
9. beta R671C 30C playback speed 3.7x 7fps
10. Emb WT 30C playback speed 3.7x 7fps
11. Emb R672C 30C playback speed 3.7x 7fps
12. Emb R672H 30C playback speed 3.7x 7fps
13. Peri WT 30C playback speed 3.7x 11fps
14. Peri R674Q 30C playback speed 4x 2fps

---

**Supplementary Figure 1.**
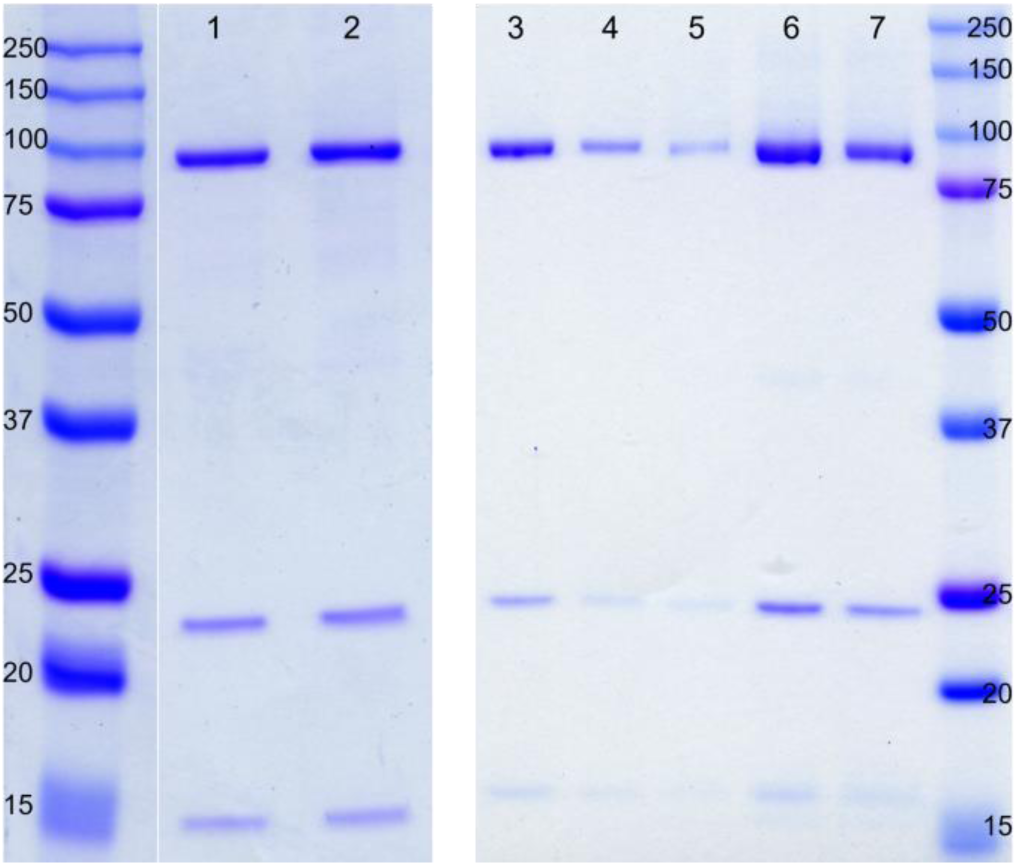
Purified myosins shown by SDS-PAGE. The purified myosin S1 trimeric complex of heavy chain, ELC, and RLC are shown for each construct in lanes 1-7. WT β (lane 1), R671C β (lane 2), WT Emb (lane 3), R672C Emb (lane 4), R672H Emb (lane 5), WT Peri (lane 6), and R674Q Peri (lane 7), and molecular weight markers labeled with corresponding kDa values.

**Supplementary Figure 2.**
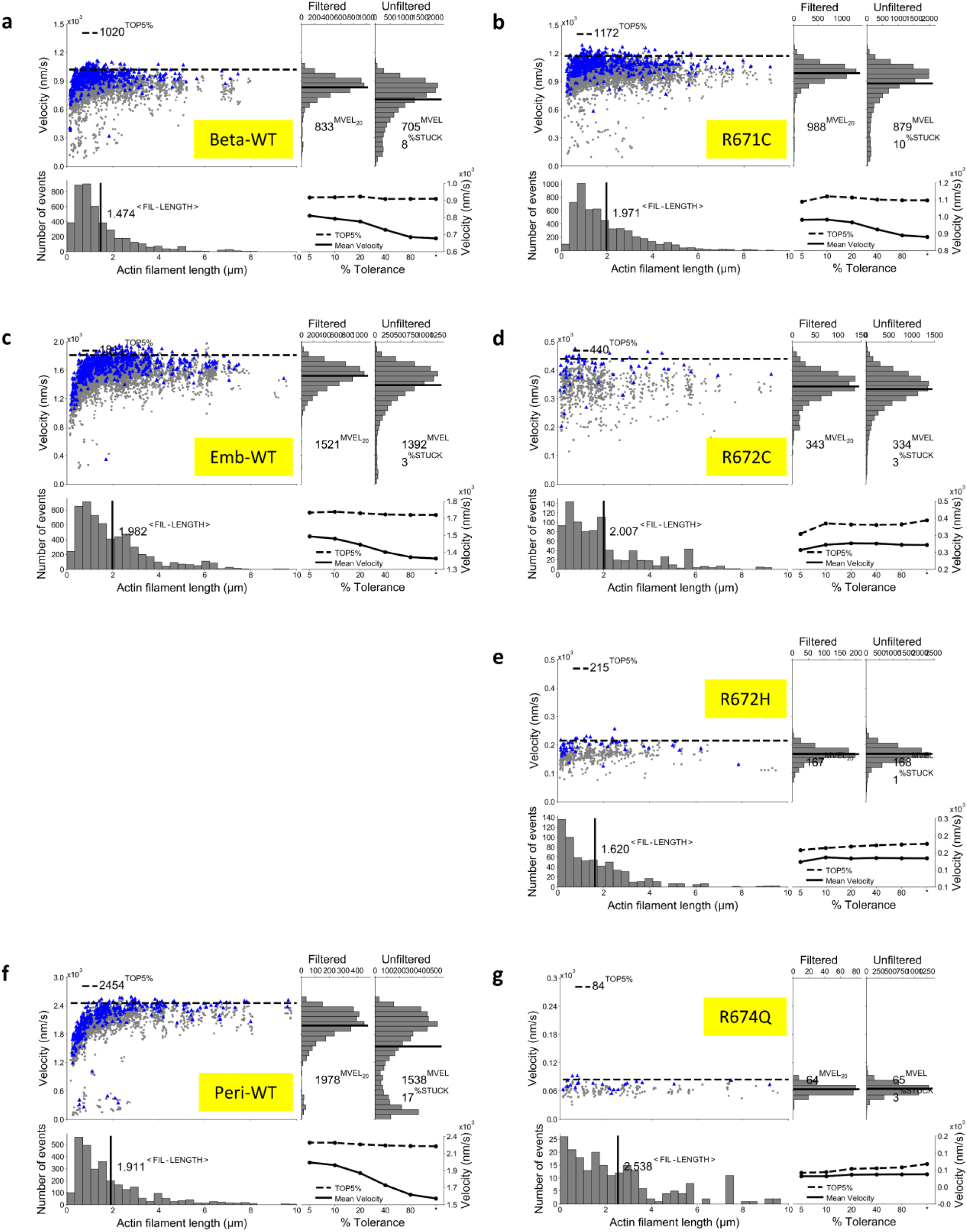
Analysis of the actin-sliding velocities of myosins in the unloaded motility assay at 23 °C by Fast Automated Spud Trekker (FAST). **a-g.** FAST analysis of data from an example motility experiment using β WT (**a**), R671C (**b**), embryonic WT (**c**), R672C (**d**), R672H (**e**), perinatal WT (**f**), and R674Q (**g**) S1. Movies from each example experiment are provided with corresponding labels in Supplementary Movies. The FAST parameters used were window size *n* = 5, path length *p* = 10, percent tolerance *pt* = 20, and minimum velocity for stuck classification *minv* = 80 nm/s (β WT and R671C) or 20 nm/s (all others). A lower *minv* was used for the embryonic and perinatal myosins to prevent possible wrongly filtering out the slow-moving filaments of their mutants. In each FAST plot, each gray data point in the top left scatter plot represents the *n*-frame averaged velocity value of a filament plotted against the filament length; thus, depending on the total number of movie frames in which a filament is present, multiple data points belong to that filament. Blue data points represent the *n*-frame averaged velocities that pass the “tolerance (*pt*)” criterion to filter out velocities with large fluctuations (see Methods for exact criteria). The “Top5%” dashed line represents the mean of the highest 5% values of the tolerance-filtered velocities. The distribution of filament lengths is shown in the bottom left histogram, with the mean shown as the vertical line. The distribution of *n*-frame averaged velocities (with and without tolerance filtering) are shown in the upper right histograms. The mean of the filtered velocities (MVEL_20_) (the metric used in the main figure in this paper, as is the convention when reporting unloaded velocities) and the mean of the unfiltered velocities (MVEL) are shown as the horizontal line in each histogram. For a filament whose velocity averaged over its entire path is less than *minv*, its *n*-frame averaged velocities are all set to 0. %STUCK is the percentage of velocities that have value 0. These 0 velocities are not shown on the FAST plots (other than via the value %STUCK) but are taken into consideration by the metric “Mean velocity including stuck (MVIS)”. Values of Top5% and MVEL_pt_ resulting from tolerance filtering at different *pt* values are given in the bottom right plots.

**Supplementary Figure 3.**
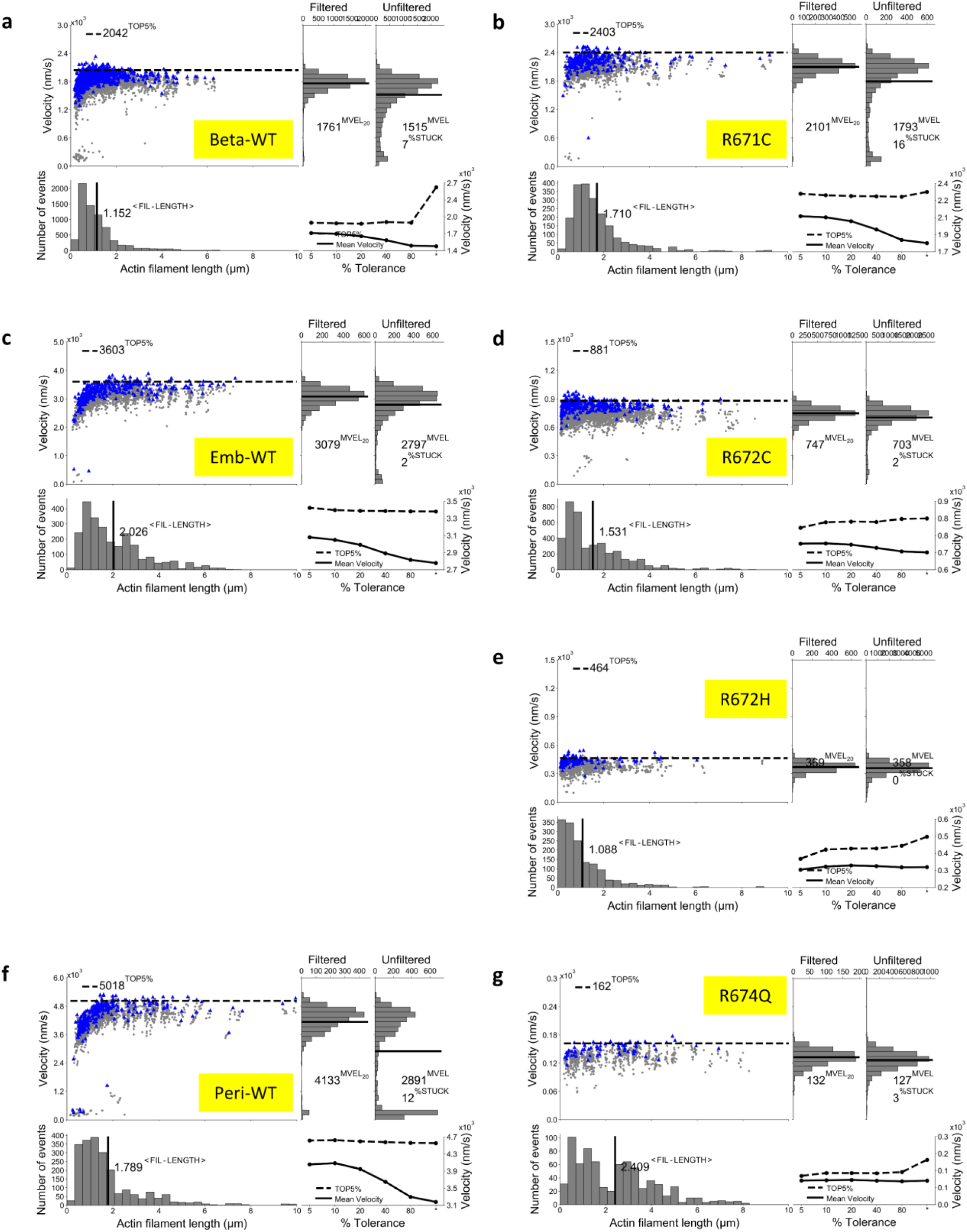
Analysis of the actin-sliding velocities of myosins in the unloaded motility assay at 30 °C by Fast Automated Spud Trekker (FAST). See Supplementary Figure 2 caption.

**Supplementary Figure 4.**
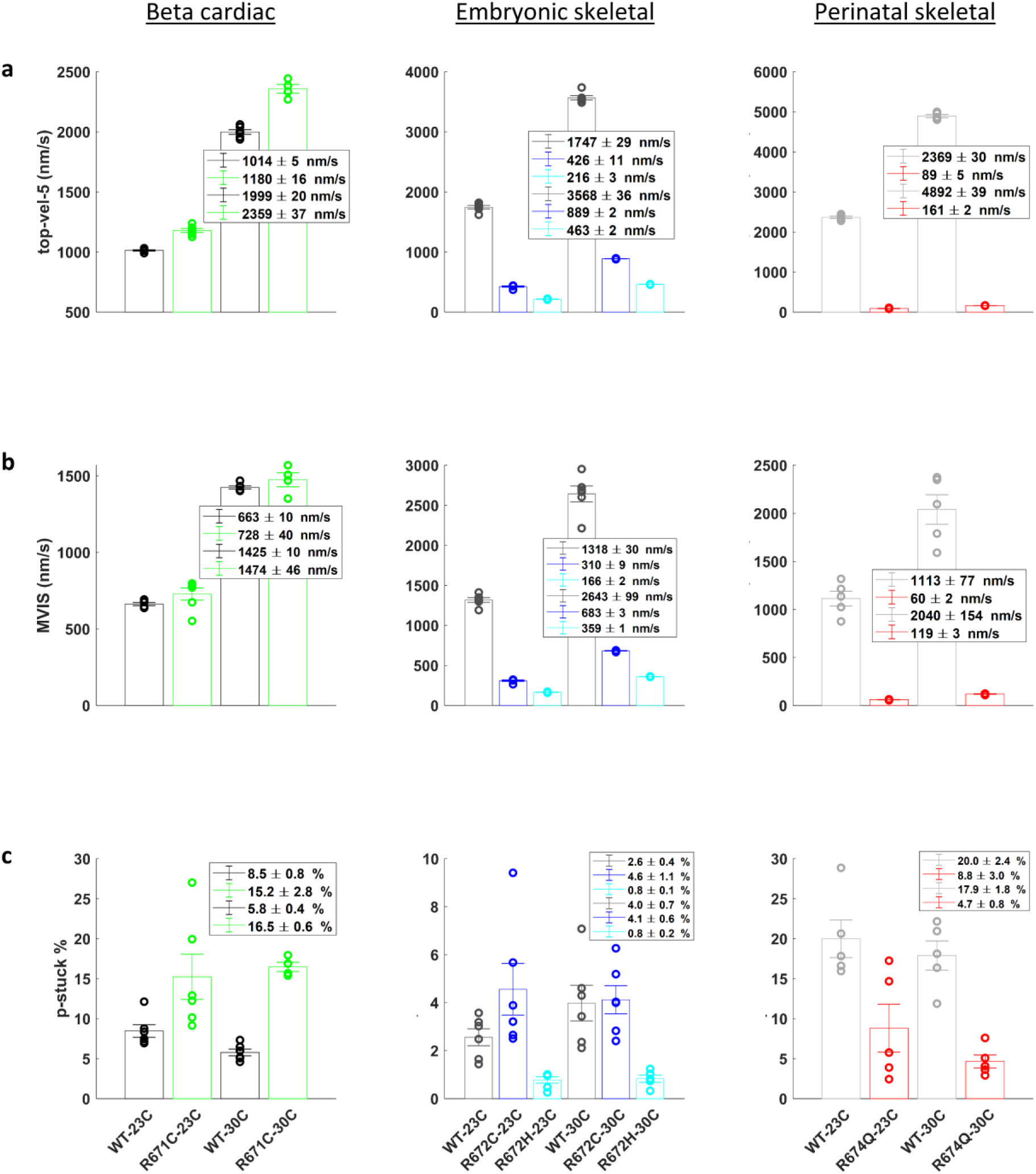
Additional motility parameters from FAST analysis. **a-c.** Top 5% velocities (**a**), mean velocity including stuck (MVIS) (**b**), and %STUCK (**c**) of all motility experiments of proteins as labeled and at 23 °C and 30 °C. Each data point represents one experiment analyzed as described in Supplementary Figures 2 and 3. Values in legends represent mean ± s.e.m.

**Supplementary Figure 5.**
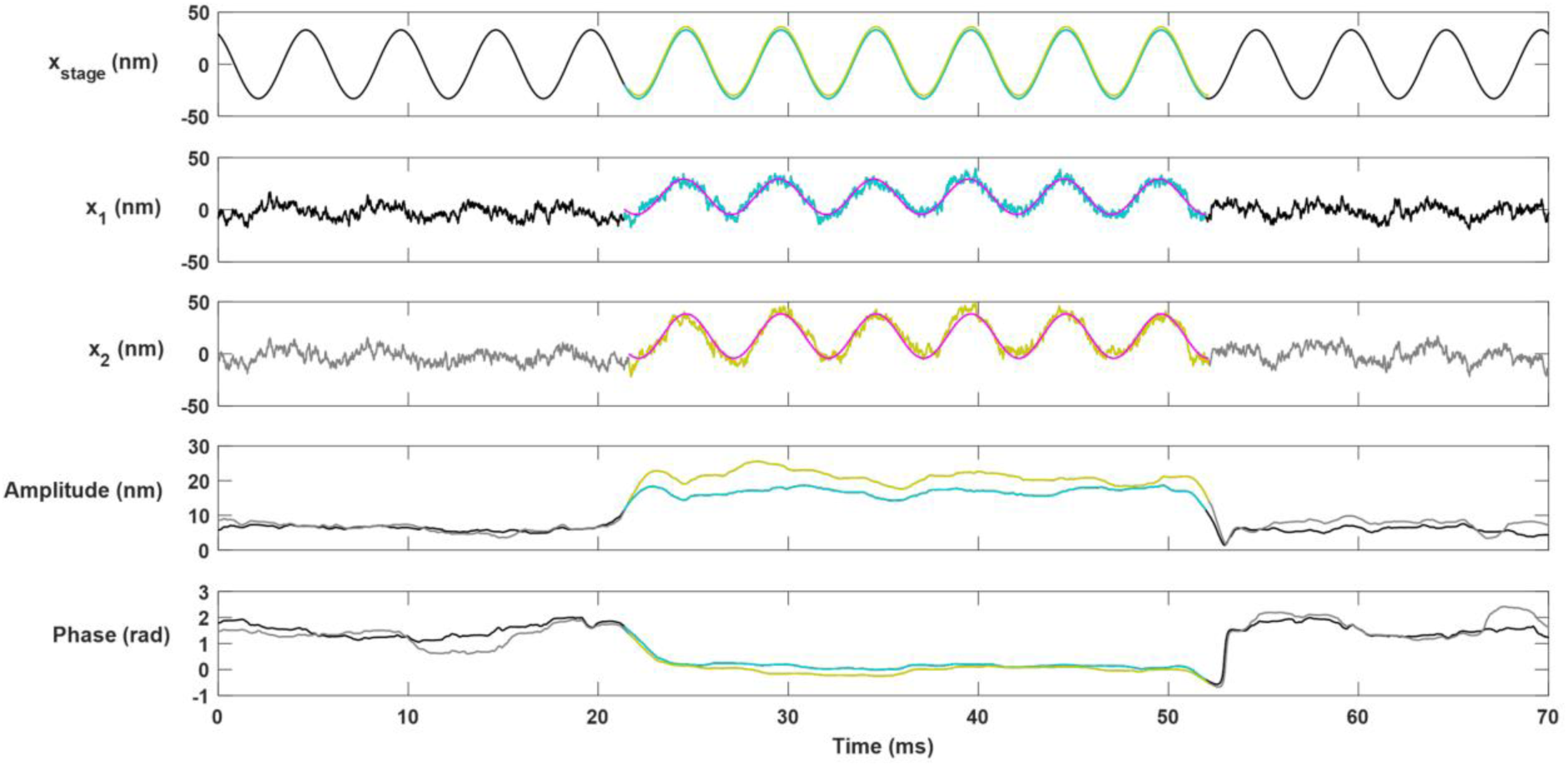
Example actin-myosin binding event from Harmonic Force Spectroscopy (HFS) optical trapping to illustrate the detection analysis method. The stage position *x*_stage_, positions of trapped beads *x*_1_ and *x*_2_, amplitudes of *x*_1_ and *x*_2_, and phases of *x*_1_ and *x*_2_ relative to the stage position are shown around one example binding event. In optical trapping using HFS, myosin sits on top of a platform bead on a stage that oscillates sinusoidally with position *x*_stage_. Upon myosin’s attachment to actin, the amplitude of the sinusoidal oscillations in *x*_1_ and *x*_2_ increases while the phase decreases because the actin dumbbell is now strongly coupled to the oscillating stage. The detected binding event is highlighted in color.

**Supplementary Figure 6.**
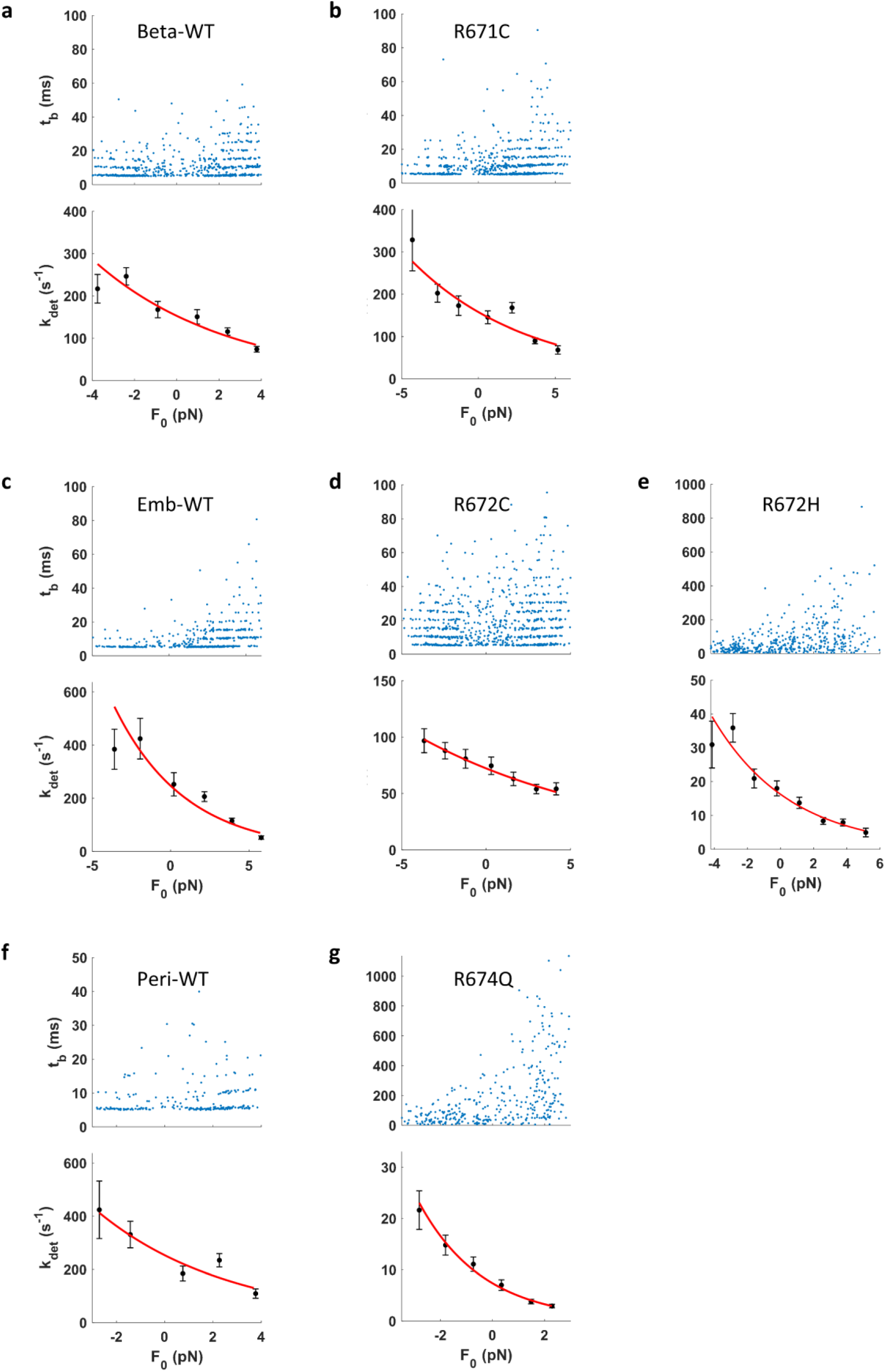
Load dependent detachment kinetics of an example molecule from each myosin measured by HFS. **a-g.** Example molecule of β WT (**a**), R671C (**b**), embryonic WT (**c**), R672C (**d**), R672H (**e**), perinatal WT (**f**), and R674Q (**g**) S1. Event lifetimes are plotted against load force (top). The detachment rate *k*_det_ at each mean force *F*_0_ is determined by maximum likelihood estimation (MLE) on the exponentially distributed lifetimes. *k*_det_ is then fitted to the Arrhenius equation with harmonic force correction (Eqn. 1) (bottom). Error bars are calculated as the variance on the MLE from the inverse Fisher information matrix (a fitting error).

**Supplementary Figure 7.**
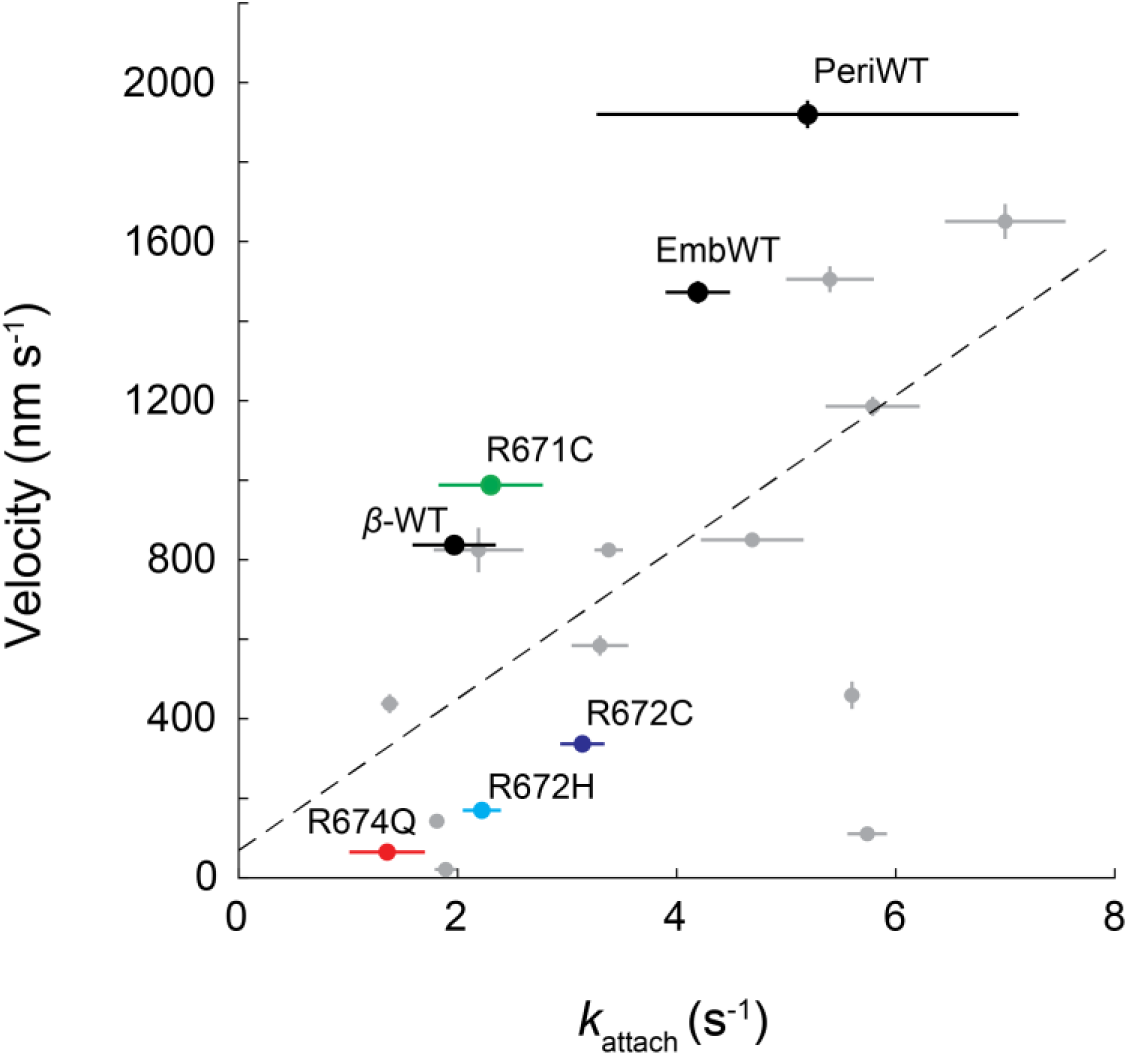
Actin sliding velocity vs. the attachment rate *k*_attach_. Linear regression gives *R*^2^ = 0.34. Gray data points are from Liu et al. *NSMB* 2018^42^. *k*_attach_ was calculated by Equation 4. Since *k*_cat_ was measured at 30 °C in the ATPase assay, while *k*_0_’s in the current paper and Liu NSMB 2018 were measured at 23 °C, we assumed *Q*_10_ = 3 for *k*_cat_ and calculated *k*_cat_’s at 23 °C to use in Equation 4 for this figure. Horizontal error bars are propagated errors from the *k*_attach_ calculation given by Equation 4. Vertical error bars are s.e.m.

**Supplementary Figure 8.**
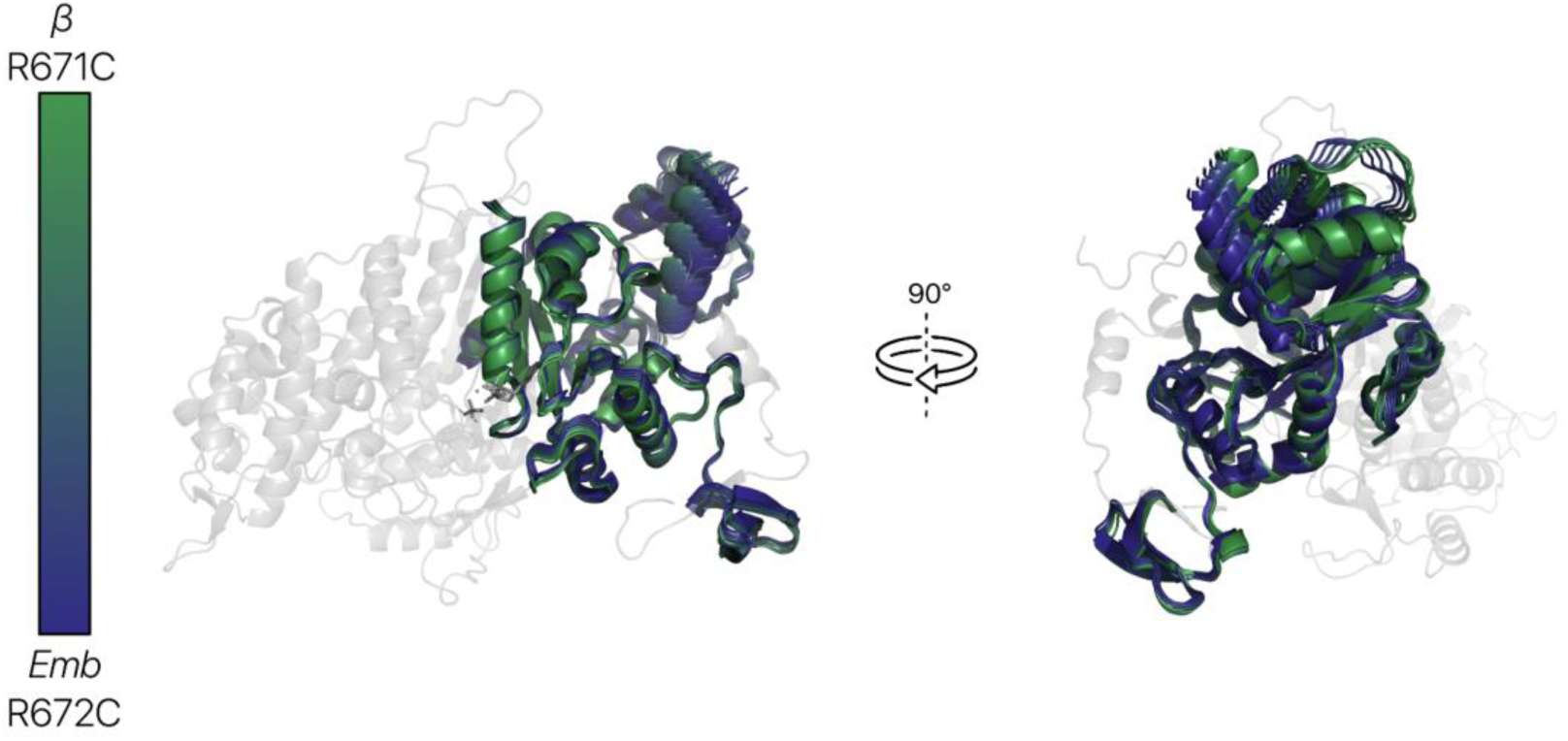
Structures generated by scanning across a DiffNets label show that lever arm down states are more probable in the R672C EmbS1 ensemble than in the R671C βS1 ensemble. Ten different structures are shown which represent the average conformation in a given DiffNets output label bin (0-0.1, 0.1-0.2, etc.). Dark blue structures are more likely to be found in the R672C EmbS1 ensemble while dark green structures are more likely to be found in the R671C βS1 ensemble.

**Supplementary Figure 9.**
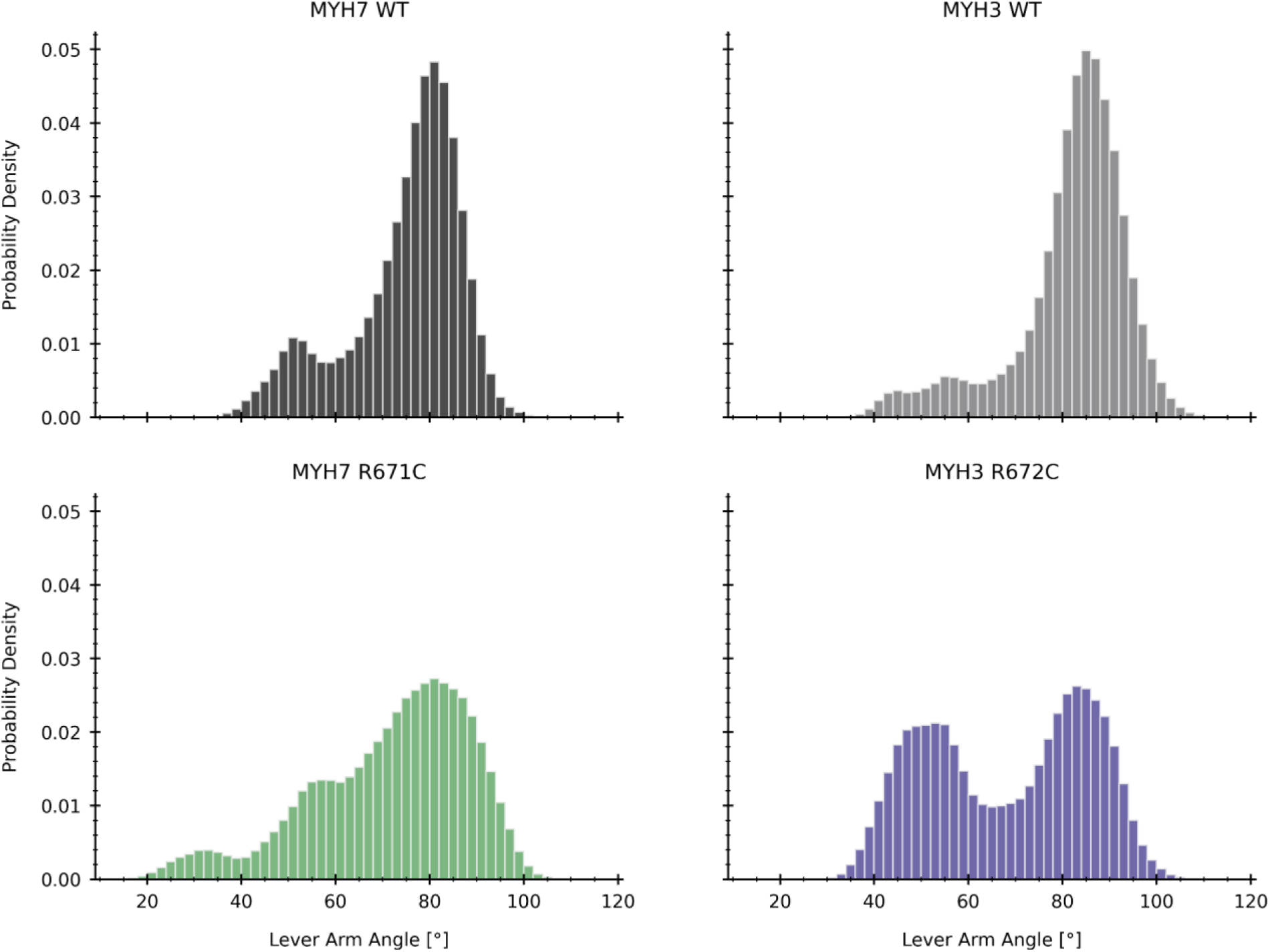
Lever arm angle distributions reveal that both embryonic R672C and β R671C are more likely to occupy lower lever arm angles but that embryonic R672C occupies lower lever arm angles (<60°) with higher probability.

**Supplementary Figure 10.**
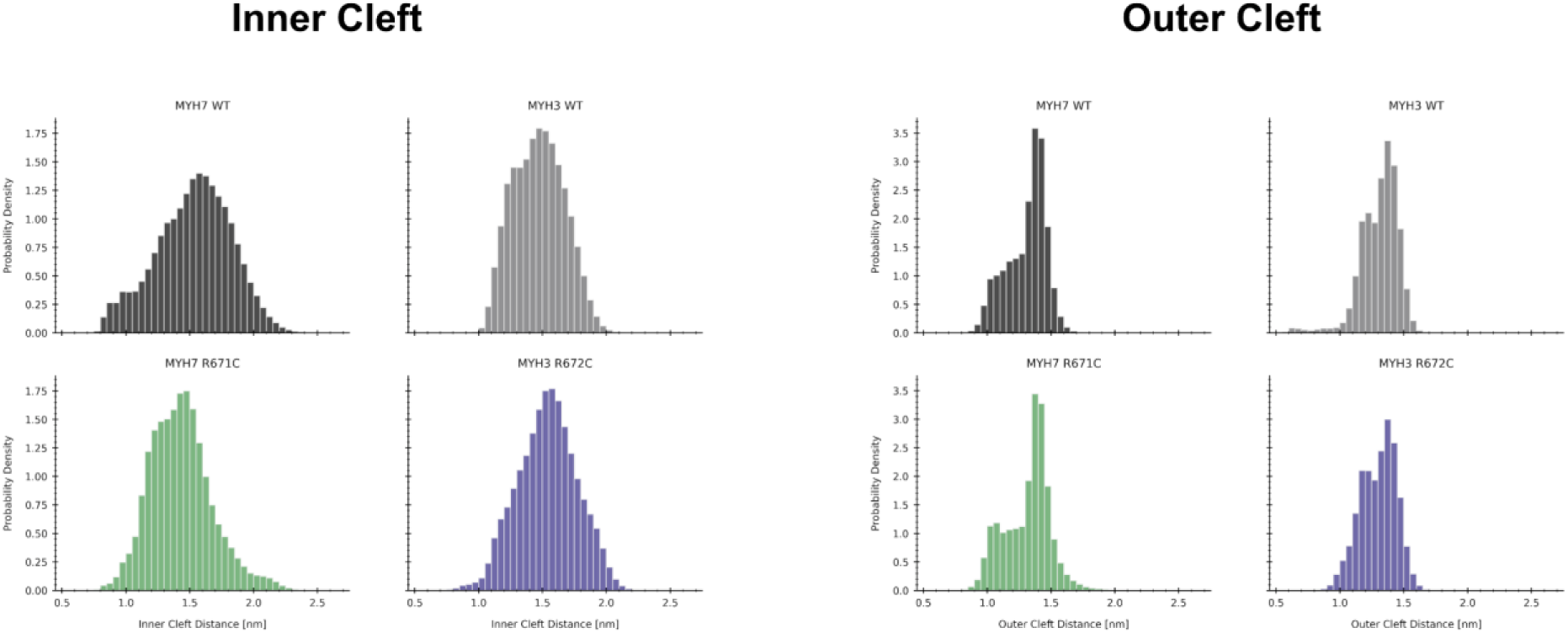
Inner and outer cleft distance distributions reveal that β myosin has a higher probability of adopting cleft-closed states in simulations. Inner cleft distances were determined based on residues L277 and S472. Outer cleft distances were determined based on residues Y422 and K598 (MYH7 numbering).

**Supplementary Figure 11.**
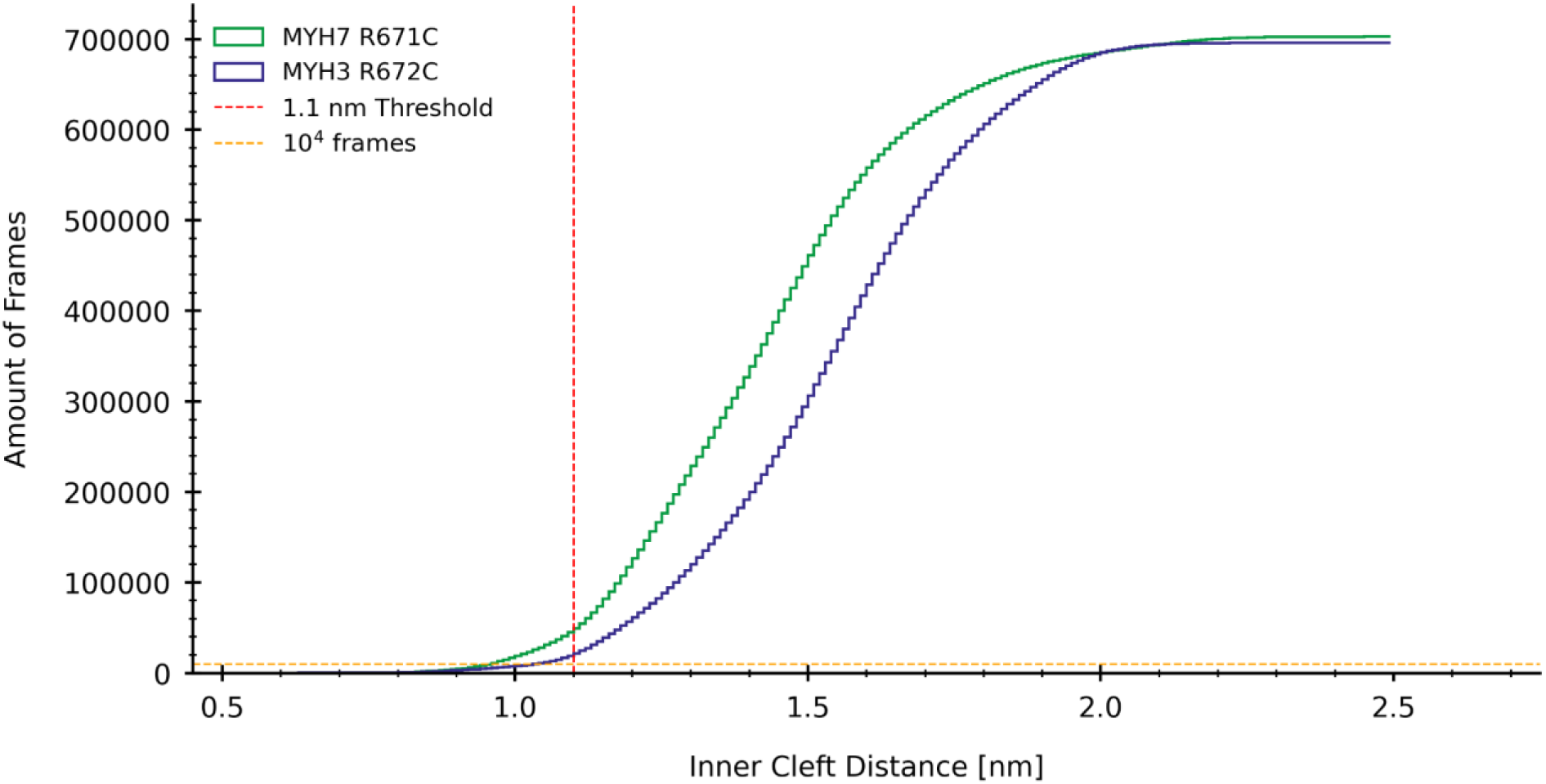
Cumulative distribution function of the inner cleft distance reveals that β R671C is more likely to occupy cleft-closed states than embryonic R672C.

**Supplementary Figure 12.**
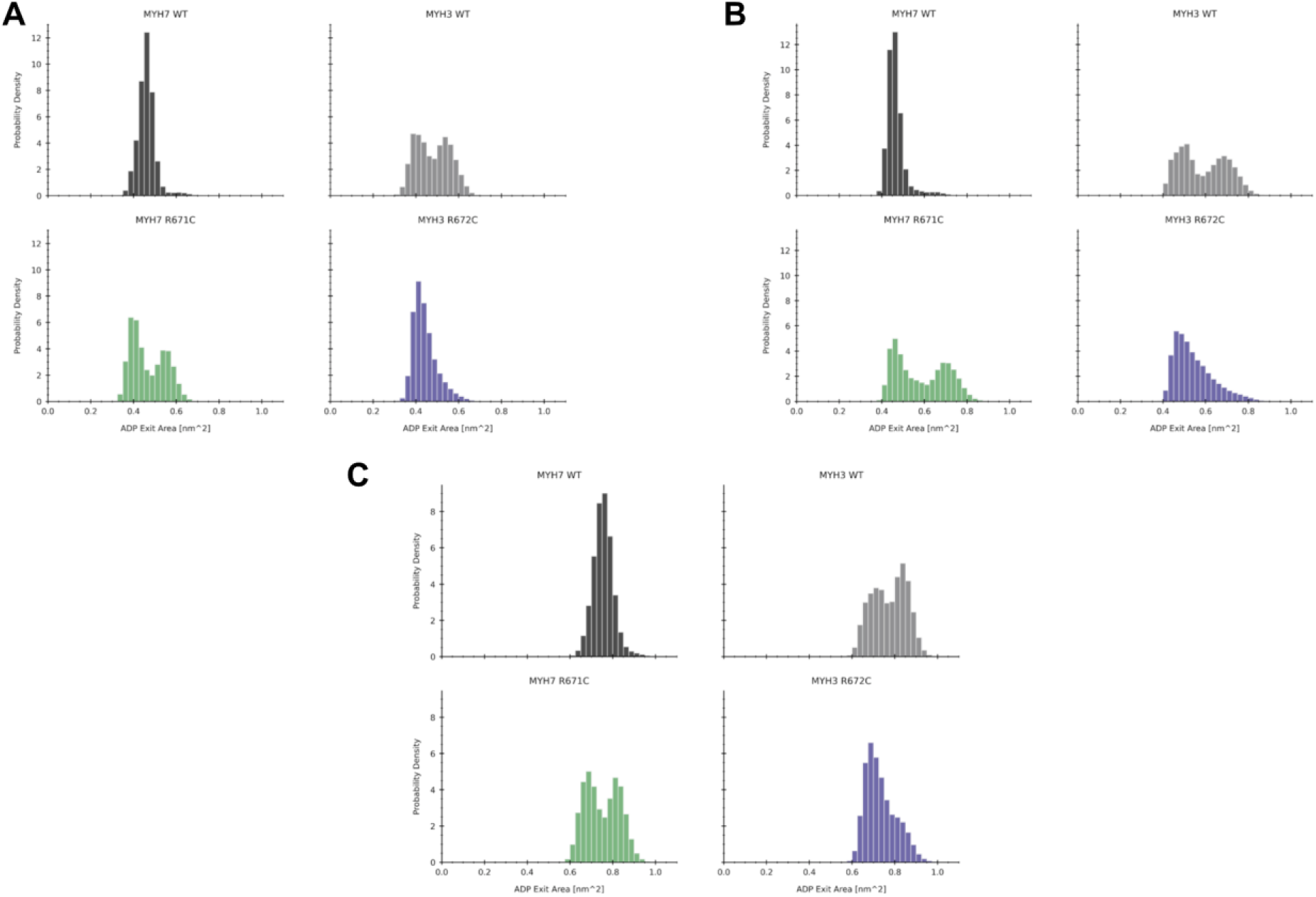
Sensitivity analysis of residues defining the ADP pocket exit area demonstrate the robustness of the method to determine ADP pocket exit area. **a.** Defining the ADP pocket exit area using the α-carbon atoms of Y128, G181, and D239 reveals that embryonic R672C, unlike the other variants, does not sample large ADP exit areas. Redefining the ADP pocket exit area using the α-carbon atoms of residues one position closer to the **(b)** N-terminus (P127, S180, N238) or **(c)** C-terminus (K129, A182, N240) maintains this relationship (MYH7 numbering).

**Supplementary Figure 13.**
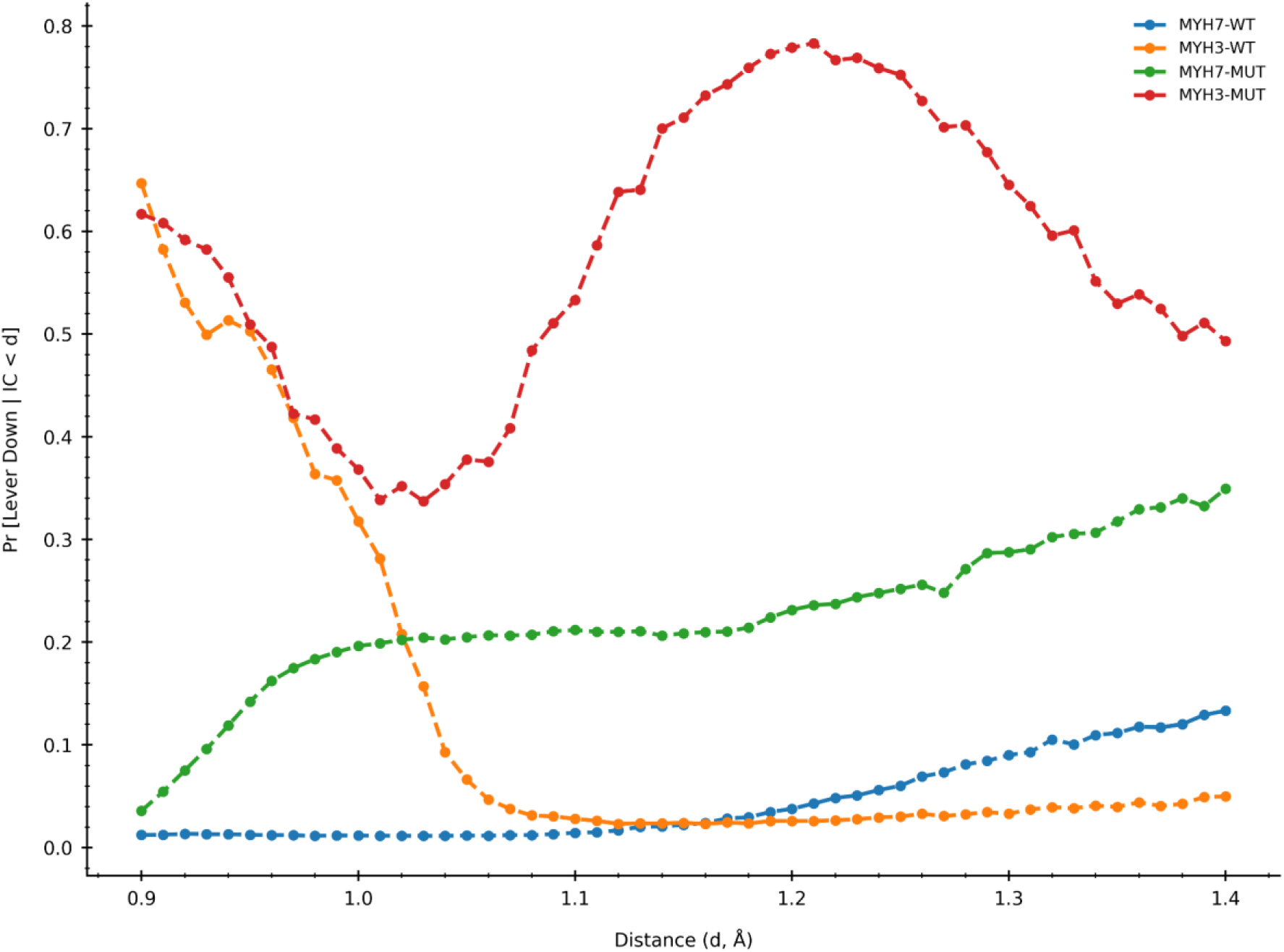
Sensitivity analysis of estimates of lever down conditional probability to inner cleft distance threshold show that embryonic R672C is more likely to occupy lever arm down states than β R671C across all inner cleft distance thresholds. A 2 Gaussian mixture model was fit to a pooled distribution of lever arm angles and each simulation frame was assigned to either the up or down Gaussian. The x-axis contains inner cleft distance thresholds that were used to condition the lever arm angle distribution. A threshold of 1.1 nm was used based on similarity to experimental actomyosin structures (reference values between 0.94 and 1.0 nm for MYH7 actomyosin structures in the PDB) and because embryonic WT contains very few frames with its lever arm closed at thresholds at or below 1 nm.

**Supplementary Table 1.** Significance tests of all measured parameters. Two sample two- sided unequal variance t-test was used. *p*-values are given. Gray numbers represent not significant (>0.05).

| Protein 1 | Protein 2 | $k_{cat}$ | $K_m$ | $V_{23C}$ | $V_{30C}$ | $k_0$ | $\delta$ | $d$ |
| --- | --- | --- | --- | --- | --- | --- | --- | --- |
| βS1 WT | EmbS1 WT | 0.012 | 0.46 | 1.8e-6 | 2.1e-8 | 0.14 | 0.011 | 0.88 |
| βS1 WT | PeriS1 WT | 0.28 | 0.14 | 5.0e-6 | 2.5e-13 | 2.2e-4 | 0.63 | 4.5e-4 |
| EmbS1 WT | PeriS1 WT | 0.64 | 0.16 | 8.2e-6 | 6.6e-8 | 3.3e-5 | 0.032 | 4.4e-3 |
| βS1 WT | βS1 R671C | 0.50 | 0.21 | 9.3e-5 | 1.2e-3 | 0.65 | 0.87 | 1.4e-3 |
| EmbS1 WT | EmbS1 R672C | 0.040 | 0.26 | 6.9e-8 | 2.0e-8 | 3.3e-4 | 2.8e-6 | 8.5e-8 |
| EmbS1 WT | EmbS1 R672H | 0.007 | 0.15 | 7.0e-8 | 1.1e-8 | 5.1e-6 | 0.013 | 5.2e-10 |
| EmbS1 R672C | EmbS1 R672H | 0.019 | 0.25 | 4.6e-8 | 9.3e-12 | 3.4e-6 | 4.6e-4 | 1.2e-3 |
| PeriS1 WT | PeriS1 R674Q | 0.20 | 0.11 | 6.6e-7 | 2.7e-9 | 5.7e-7 | 8.5e-4 | 6.4e-7 |

